# Metagenomics-enabled proteomics reveals how AMF and PSB co-inoculation reshapes tomato rhizosphere dynamics across growth stages

**DOI:** 10.64898/2026.05.12.724390

**Authors:** Yejin Son, Eric J. Craft, Miguel A. Pineros, Olivia L. Mathieson, Ayesha Awan, J. Alfredo Blakeley-Ruiz, Manuel Kleiner, Jenny Kao-Kniffin

## Abstract

Urban agriculture increasingly relies on compost-based substrates for sustainable production, yet we lack a clear characterization of how these systems respond to biological amendments aimed at introducing beneficial microbiota. Here we investigated how developmental stage and co-inoculation with arbuscular mycorrhizal fungi (AMF) and phosphate-solubilizing bacteria (PSB) reshape rhizosphere microbial function in *Solanum lycopersicum* grown in compost-based urban farm substrate. Using plant physiology assays, 16S rRNA amplicon sequencing, and metagenome-informed metaproteomics, we characterized tomato physiological responses and rhizosphere microbial activity during flowering and fruiting across control, single AMF, single PSB, and AMF and PSB co-inoculation treatments. Co-inoculation synergistically enriched beneficial taxa, improved fruit nutrient accumulation, elevated nutrient transporter and quorum sensing protein production, and drove stress-driven dormancy in competitively excluded taxa, with responses varying between developmental stages. Our findings establish metagenome-informed metaproteomics as essential for resolving stage-specific rhizosphere microbiome functional responses to tomato development and AMF and PSB co-inoculation.

## Introduction

While beneficial bacteria and fungi are commonly utilized in plant production systems, the synergistic effects of combining multiple microbial taxa remain largely unexplored^1^.

Microbial communities are well known to drive soil nutrient cycling by enzymatically decomposing organic matter and mineralizing key elements such as P and N into plant available forms^2^. Among these consortia, the synergistic partnership among plants, arbuscular mycorrhizal fungi (AMF), and phosphate solubilizing bacteria (PSB) is particularly effective in promoting plant growth and improving soil fertility. AMF have co evolved with plants for over 400 million years and associate with most terrestrial species, exchanging mineral nutrients for plant derived carbon and channeling photosynthetically fixed carbon into the rhizosphere, where it fuels beneficial bacteria such as PSB^3–7^. These bacteria reciprocate by decomposing urea and organic P, enhancing nutrient availability and reinforcing a positive feedback loop that supports AMF and plant nutrition and microbial activity. Empirical studies show that AMF and PSB co inoculation improves tomato growth and P uptake in phospho compost systems^8^ and enhances Ca, K, and P uptake in green waste composts under drought stress^9^. Together, these findings highlight the potential of microbial consortia to improve nutrient use efficiency, stabilize nutrient cycling, and reduce nutrient losses in compost-based systems.

Despite clear evidence that AMF and PSB co inoculation enhances plant nutrient uptake, growth, and stress resilience^8,9^, how AMF, PSB, and their combined application influence compost microbiome structure and function remains largely unknown. Previous studies using16S rRNA gene profiling or metagenomics have shown that microbial inoculants can alter native soil microbiota composition, but these approaches offer limited insight into underlying mechanisms because they provide only coarse taxonomic resolution and cannot directly assess microbiome function^10,11^. Integrating metagenomics with metaproteomics offers a powerful means to profile microbial proteins and reveal physiological responses to inoculation, and it also expands the resolution at which soil microbiome function can be interpreted. Building on these methodological advances, the present study investigates how AMF, PSB, and their co inoculation influence rhizosphere microbial functions across flowering and fruiting stages of tomato plants grown in constructed soils. To address this gap, we conducted a controlled growth chamber experiment using tomato plants subjected to four inoculation treatments: a non-inoculated control group, inoculation with AMF only (AMF tomatoes), inoculation with PSB only (PSB tomatoes), and co-inoculation with AMF and PSB (hereafter referred to as AMF+PSB tomatoes). We hypothesized that co-inoculation with AMF and PSB would result in a synergistic growth response in tomato plants, outperforming both single-taxon treatments and the non-inoculated control. By combining the phosphorus-solubilizing capabilities of PSB with the expansive transport network of AMF, we expect a significant increase in phosphorus accumulation within plant tissues and fruit, ultimately driving greater tomato biomass.

## Results

We evaluated microbially-mediated changes in the physiology and nutrient status of tomato plants at flowering and fruiting stages, in addition to measuring AMF colonization, microbiome profiling (through 16S rRNA amplicon sequencing and metagenomics) and microbiome function (through metaproteomics and soil enzymatic activity) (Figure 1). Metagenomic sequencing was performed on triplicate soil samples from control and AMF+PSB tomatoes at each developmental stage, as the co-inoculated metagenome encompassed organisms from both single-inoculation treatments. Shallow sequencing of individual replicates generated an average of ∼71 million paired-end reads per sample, while deep sequencing of pooled samples yielded an average of 541 million paired-end reads. Co-assembly and binning produced 1,058 high to medium quality metagenome-assembled genomes (MAGs) (≥ 50% completeness, < 10% contamination)^12^. Metaproteomic analysis of all 72 rhizosphere samples (9 replicates per condition) by LC-MS/MS identified 26,961 protein groups, together providing functional insights into microbial activity and interactions at the tomato root-soil interface.

**Figure 1.**
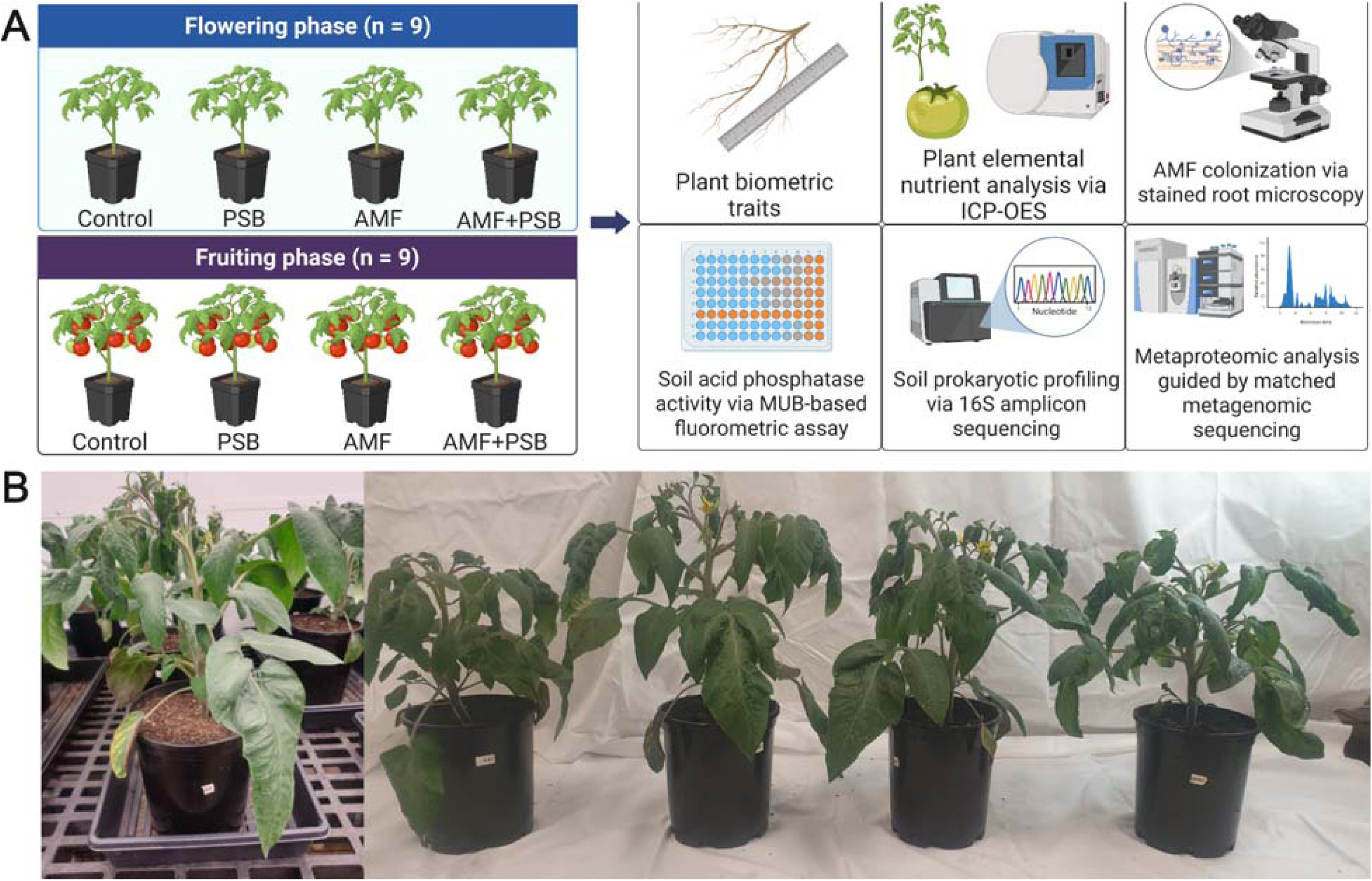
Overview of experimental design and methodologies. (A) Schematic representation of tomato experiments conducted at two developmental stages (flowering and fruiting) to assess treatment effects. Treatment groups included and uninoculated controls, PSB (plants inoculated with phosphate solubilizing bacteria), AMF (plants inoculated with arbuscular mycorrhizal fungi), and AMF+PSB (plants co inoculated with both). (B) Representative images of tomato plants during the flowering stage.

### Co-inoculation with AMF and PSB leads to strong impacts on plant phenotype and nutrient content

Co-inoculation of AMF and PSB proved highly effective in improving tomato growth, nutrient absorption, and mycorrhizal colonization throughout plant development. At fruiting, co-inoculated tomatoes outperformed controls in shoot height, dry mass, and root length, confirming a synergistic effect on plant biomass (Figure 2A). AMF alone raised Fe, K, P, and S levels in shoot tissues, while co-inoculation further enhanced Mg and Zn accumulation. Fruits from co-inoculated plants contained higher concentrations of Fe, K, Mg, P, S, and Zn than controls, with reduced shoot K alongside elevated fruit K suggesting enhanced K translocation (Figure 2B). At flowering, co-inoculated plants showed markedly higher AMF colonization than AMF-only plants, indicating PSB-mediated enhancement of early mycorrhizal establishment, though this advantage diminished at fruiting (Figure 2C). Low but non zero colonization in control and PSB tomatoes reflected background AMF propagules in the compost, although the introduced inoculants were substantially more effective. Microscopic images of AMF colonization structures in tomato roots are provided in Supplementary Figure 1. Acid phosphatase activity followed a similar pattern, with AMF+PSB tomatoes showing the highest activity at flowering, followed by AMF and PSB tomatoes, all exceeding controls (Figure 2C).

**Figure 2.**
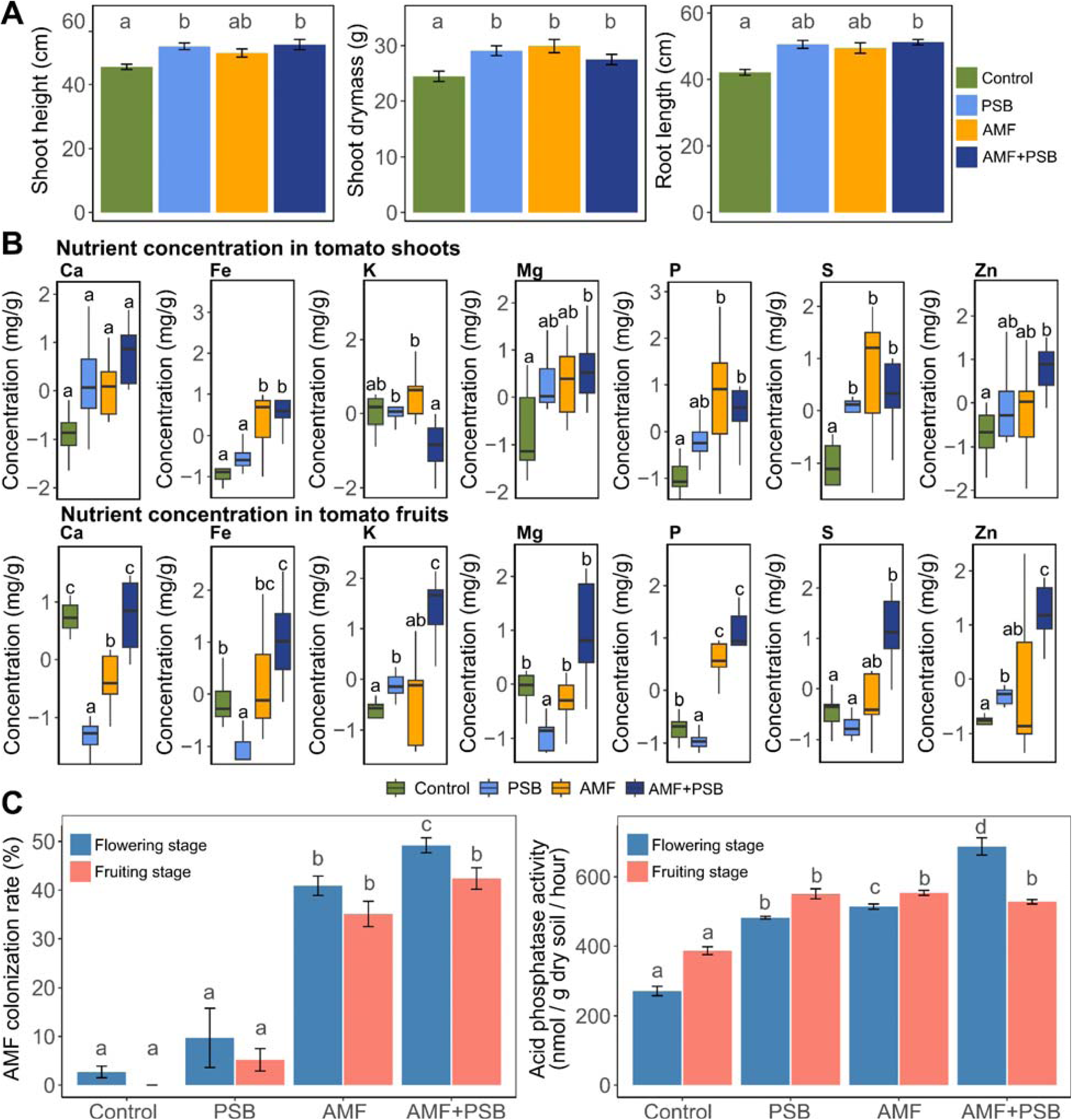
Comparative analysis of tomato growth parameters, nutrient composition, AMF colonization rates, and soil acid phosphatase activity. (A) Growth traits, including shoot height, shoot dry mass, and root length were measured in ten-week-old tomato plants at the fruiting stage (n = 9). The X-axis represents the various microbial treatments applied. Values are reported as means ± standard error, with statistical significance determined using pairwise Wilcoxon rank-sum tests at Benjamini-Hochberg (BH) adjusted *p* values < 0.05. (B) Box plots displaying nutrient concentrations in tomato shoots and fruits at the fruiting stage, quantified via inductively coupled plasma optical emission spectroscopy (ICP-OES) (n = 9). Box colors indicate treatment categories, and statistical significance within each nutrient type across treatments is evaluated using pairwise Wilcoxon rank-sum tests at BH adjusted *p* < 0.05. (C) AMF colonization rates and soil acid phosphatase activity are presented for both flowering and fruiting stages (n = 9). AMF colonization was verified by ink-vinegar staining, and acid phosphatase activity was quantified using a fluorometric enzyme assay with 4-methylumbelliferyl phosphate as the substrate. Significant differences within each plant developmental stage are assessed using pairwise Wilcoxon rank-sum tests with BH adjusted *p* values < 0.05. Statistical significance is denoted by distinct alphabetical letters, where different letters indicate significantly different groups. Abbreviations: PSB, tomatoes treated with phosphate-solubilizing bacteria (PSB) only; AMF, tomatoes treated with arbuscular mycorrhizal fungi (AMF) only; AMF+PSB, tomatoes co-inoculated with both AMF and PSB.

### Microbial inoculation strongly impacts Rhizosphere Prokaryotic Community Structure and Composition

The diversity decline accompanying the transition to fruiting was most pronounced in control and PSB tomatoes, suggesting AMF-based inoculations better sustained community richness through plant development. Pairwise PERMANOVA confirmed significant compositional differences across both stages and treatments (Figure 3A; Supplementary Table 1), with non-overlapping confidence ellipses for AMF and AMF+PSB relative to controls at flowering, and further divergence by fruiting where all treatments formed distinct clusters. Rhizosphere microbial diversity fluctuated across treatments and stages, with PSB tomatoes recording the greatest diversity at flowering, while AMF and AMF+PSB tomatoes had significantly higher diversity than controls and PSB at fruiting (Figure 3B).

**Figure 3.**
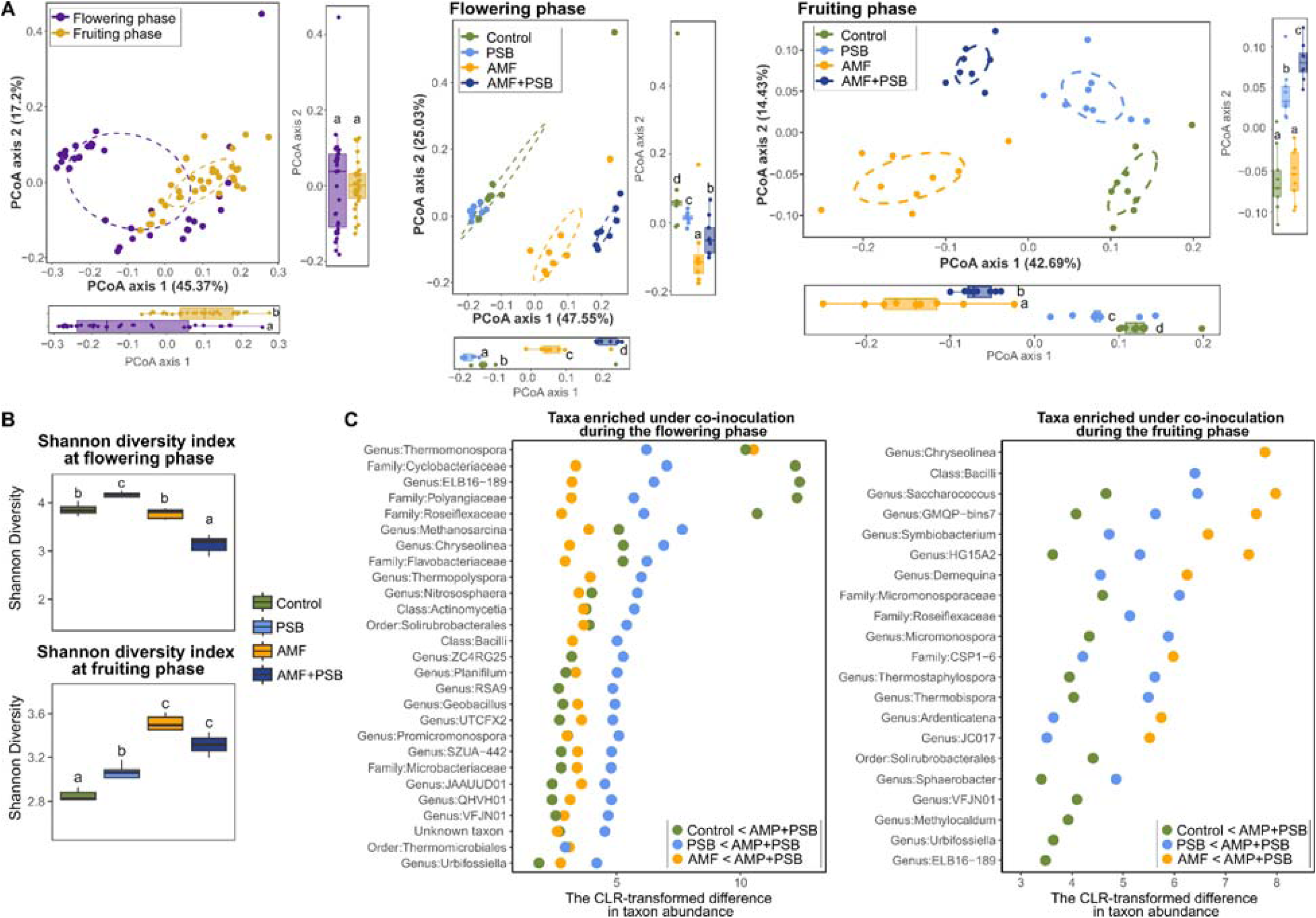
Microbial diversity patterns and differentially abundant taxa across inoculation treatments, based on 16S rRNA gene amplicon sequencing data. (A) Principal coordinates analysis (PCoA) of soil prokaryotic communities, based on Bray-Curtis dissimilarities calculated from genus level relative abundances derived from 16S rRNA gene amplicon sequencing, to visualize developmental shifts and treatment driven differences in community structure. The 95% confidence ellipses for each site are drawn to represent treatment-specific clustering. Additionally, box plots for PCoA axis 1 and PCoA axis 2 values are displayed at the bottom and right of each PCoA plot, providing further insights into clustering patterns. Statistical significance of differences along PCoA axes was determined using the pairwise Wilcoxon rank-sum test at BH adjusted *p* < 0.05, with different alphabetic letters denoting significant groupings. (B) Shannon diversity indices of microbial communities corresponding to each inoculation treatment at the flowering and fruiting developmental stages of tomato plants. Shannon index values were calculated from genus-level relative abundance profiles and compared across treatments using the pairwise Wilcoxon rank-sum test at Benjamini-Hochberg (BH) adjusted *p* < 0.05, with significance denoted by different alphabetic letters. (C) Enriched microbial taxa under co-inoculated conditions identified by differential abundance analysis using ALDEx2-based differential abundance analysis of genus level profiles obtained during the flowering and fruiting stages of tomato development^13^. Centered log-ratio (CLR) scores reflect treatment-driven differences in taxon abundance, with positive values indicating enrichment and negative values indicating relative depletion in co-inoculated tomatoes; only taxa significantly enriched in the AMF+PSB treatment compared with the control, PSB, and AMF groups are displayed, while Supplementary Figure 2 includes both enriched and depleted taxa. The Wilcoxon rank-sum test was applied to identify significantly enriched microbial taxa, with *p* values corrected using the BH method (*p* < 0.05). Each circle is color-coded to represent different comparison approaches. Abbreviations: Control, uninoculated tomatoes; PSB, treated with phosphate-solubilizing bacteria only; AMF, treated with arbuscular mycorrhizal fungi only; AMF+PSB, co-inoculated with both AMF and PSB.

Prior to differential abundance analysis with ALDEx2^13^, we applied an 70% prevalence filter, retaining 101 taxa for subsequent testing across treatments, whose CLR-transformed counts across microbial treatments and developmental stages are summarized in Supplementary Table 2. Of the tested taxa, 83 showed significant differential abundance among treatments, with Supplementary Table 3 reporting their CLR differences, median effect sizes, and adjusted p values. These taxa encompassed groups associated with co inoculation as well as those enriched or depleted in the control, AMF, and PSB treatments. Of these, 65 taxa were associated with co inoculation, showing either increased or decreased abundance relative to the control, AMF, and PSB treatments (Supplementary Figure 2). Figure 3C specifically depicts the taxa enriched in the rhizosphere of co inoculated tomato plants during both the flowering and fruiting phases. During the flowering stage, enrichment patterns included members of the classes Actinomycetia and Bacilli; families Cyclobacteriaceae, Polyangiaceae, Roseiflexaceae, Flavobacteriaceae, and Microbacteriaceae; and a broad set of genera such as *Thermomonospora*, ELB16-189, *Methanosarcina*, *Chryseolinea*, *Thermopolyspora*, *Nitrososphaera*, ZC4RG25, *Planifilum*, RSA9, *Geobacillus*, UTCFX2, *Promicromonospora*, SZUA-442, JAAUUD01, QHVH01, VFJN01, and *Urbifossiella*, along with the orders Solirubrobacterales and Thermomicrobiales. During the fruiting stage, enriched taxa included the class Bacilli; families Micromonosporaceae, Roseiflexaceae, and CSP1-6; and several genera including *Chryseolinea*, *Saccharococcus*, GMQP-bins7, *Symbiobacterium*, HG15A2, *Demequina*, *Micromonospora*, *Thermostaphylospora*, *Ardenticatena*, JC017, *Sphaerobacter*, VFJN01, *Methylocaldum*, *Urbifossiella*, and ELB16-189, along with the order Solirubrobacterales.

### Genome-Resolved Insights into Rhizosphere Microbial Shifts Induced by AMF and PSB Co-Inoculation Across Tomato Developmental Stages

MAG analysis (Figure 4A), restricted to MAGs with ≥ 50% completeness and < 10% contamination^12^ from the triplicate shallow-sequencing dataset, revealed a phylogenetically diverse rhizosphere community dominated by Pseudomonadota (200), Actinomycetota (161), Planctomycetota (94), Chloroflexota (80), Bacteroidota (84), Gemmatimonadota (66), and Myxococcota (71), with 118 additional MAGs distributed across Bacillota subgroups (Supplementary Table 4). Family-level comparison between MAG and 16S rRNA gene ASV datasets identified 9 overlapping families with relative abundance greater than 1%, including Cyclobacteriaceae, Hyphomicrobiaceae, Microbacteriaceae, Micromonosporaceae, Polyangiaceae, RSA9, Steroidobacteraceae, Streptosporangiaceae, and UBA4823 (Figure 4B). At the broader family level, the two methods shared 78 families (21% of the total), while MAGs exclusively included an additional 273 families, highlighting the greater taxonomic depth offered by metagenomic sequencing (Figure 4C). Metagenomic profiling identified 130 unique MAGs at the species level, 14 of which exceeded 0.5% relative abundance (Figure 4D), encompassing both characterized species (*Methylocaldum szegediense*, *Mycobacterium hassiacum*, and *Sphaerobacter thermophilus*) and uncultured or candidate lineages (JAAYXE01 sp012516075, RSA1 sp002919305, and multiple ZC4RG lineages).

**Figure 4.**
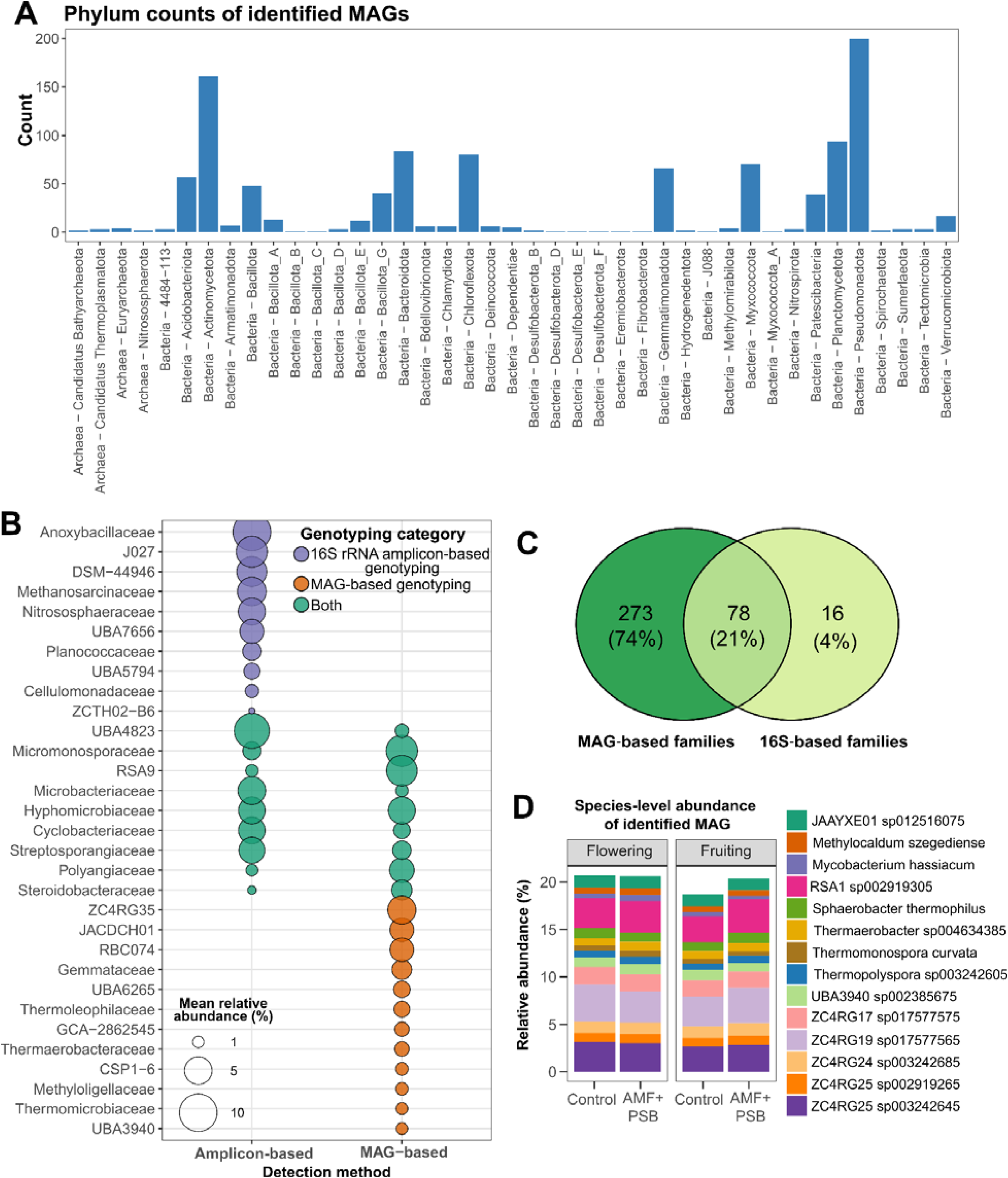
Taxonomic composition of the tomato rhizosphere during flowering and fruiting stages based on prokaryotic metagenome-assembled genome (MAG) abundances. (A) Distribution of prokaryotic MAG counts across microbial phyla, comprising dereplicated high– to medium-quality MAGs with ≥ 50% completeness and < 10% contamination. Taxonomic hierarchy is denoted as kingdom-phyla to ensure clear differentiation of broad classifications. (B) Cross-method comparison of family-level taxonomic abundance profiles derived from 16S rRNA gene amplicon sequencing (ASV-based, n = 9) and MAG-based shallow sequencing (n = 3), restricted to families exceeding 1% median relative abundance in at least one method. Bubble size reflects mean relative abundance within each detection method. Families exclusive to 16S rRNA amplicon sequencing are shown in purple, MAG-exclusive families in red, and those detected by both sequencing approaches in green. (C) Venn diagram comparing family-level taxonomic overlap between MAG-based and 16S rRNA gene ASV-based approaches, showing the number and percentage of shared and method-exclusive families detected across all samples. (D) Species-resolved MAG abundances derived from shallow metagenomic sequencing of triplicate samples (n = 3) captured taxa exceeding 0.5% relative abundance. Abbreviations: Control, uninoculated tomatoes; PSB, treated with phosphate-solubilizing bacteria only; AMF, treated with arbuscular mycorrhizal fungi only; AMF+PSB, co-inoculated with both AMF and PSB.

### Differences in Tomato Rhizosphere Metaproteome across Microbial Inoculation Treatments and Plant Developmental Stages

The metaproteome showed significant compositional differences across both tomato developmental stages and microbial treatments, as supported by PCA clustering (Figure 5A) and pairwise PERMANOVA (Supplementary Table 1). At flowering, the comparable metaproteomic profiles of AMF and AMF+PSB tomatoes indicated that AMF was the primary driver of protein abundance patterns; by fruiting, their divergence revealed a progressively stronger co-inoculation effect as plants matured (Supplementary Figure 3; Supplementary Table 5). The major KEGG Level 3 pathways enriched under co-inoculation, included quorum sensing (QS), ABC transporters, cell growth, and transport (Figure 5B, Supplementary Table 6), with taxonomic contributors detailed in Figure 5C, and (Supplementary Table 7).

**Figure 5.**
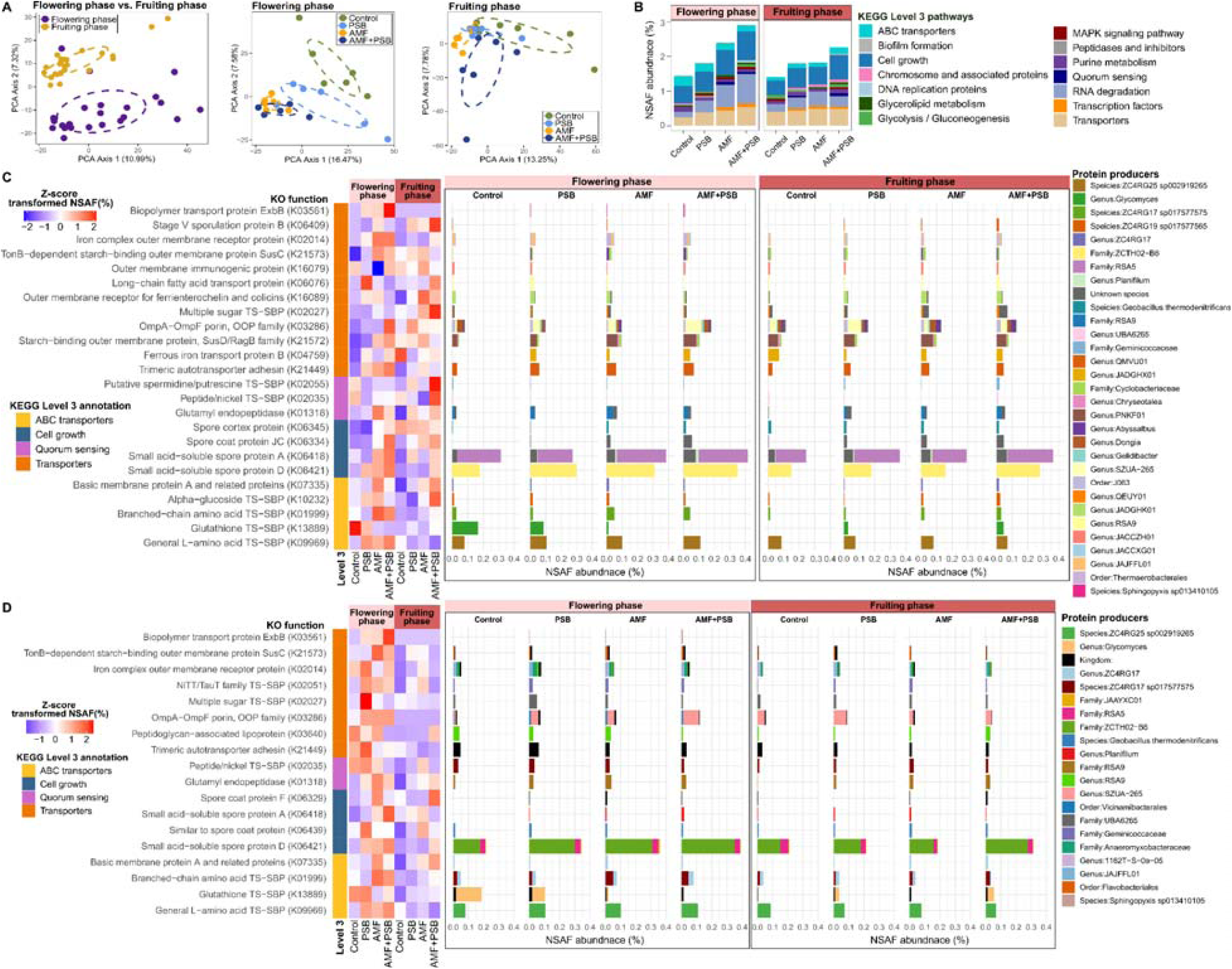
Functional shifts in the tomato rhizosphere metaproteome under microbial inoculation across developmental stages. (A) PCA of CLR-transformed metaproteomic profiles across inoculation treatments at flowering and fruiting stages, with 95% confidence ellipses denoting treatment-level clustering. (B) Cumulative normalized spectral abundance factor (NSAF) mean abundances of individual proteins significantly enriched under AMF and PSB co-inoculation relative to controls and single-inoculation treatments, aggregated by KEGG Level 3 pathway and color-coded by functional category (n = 6). Differential enrichment was assessed at the individual protein level by Welch’s two-sample t-test with Benjamini-Hochberg (BH) correction (log□ fold change > 0.5, adjusted *p* value < 0.05), and only KEGG Level 3 pathways exceeding 0.05% cumulative NSAF abundance are shown. (C) Heatmap of microbial proteins significantly enriched under AMF and PSB co-inoculation relative to control and single-inoculation treatments (Welch’s two-sample t-test, BH adjusted *p* < 0.05, log□ fold change > 0.5), restricted to KEGG Level 3 categories of transporter activity, cell growth, and quorum sensing. (D) Heatmap of microbial proteins significantly enriched during flowering relative to fruiting within each inoculation treatment (Control, PSB, AMF, and AMF+PSB), identified by Welch’s two-sample t-tests with Benjamini-Hochberg (BH) adjusted *p* < 0.05 and log□ fold change > 0.5, restricted to KEGG Level 3 categories of transporter activity, cell growth, and quorum sensing. Rows represent individual proteins grouped by KO function and annotated by KEGG Level 3 category, with heatmap colors reflecting Z-score-standardized NSAF values (red, high abundance; blue, low abundance). The eight columns represent four inoculation treatments (Control, PSB, AMF, and AMF+PSB) at two developmental stages, with flowering (columns 1-4) and fruiting (columns 5-8) shown left to right. Each horizontally stacked bar displays the mean NSAF abundance (%) of a given enriched protein, with individual segments representing the contribution of each microbial taxon, color-coded by microbial source as indicated in the legend, across four inoculation treatments (Control, PSB, AMF, and AMF+PSB; n = 6 replicates per treatment) at two developmental stages (flowering and fruiting). *Abbreviations:* Control, uninoculated tomatoes; PSB, phosphate-solubilizing bacteria only; AMF, arbuscular mycorrhizal fungi only; AMF+PSB, co-inoculated with both; TS-SBP, transport system substrate-binding protein.

During flowering, co-inoculated tomatoes showed greater abundance of several proteins relative to control and single-inoculation treatments (Figure 5C), spanning nutrient transport, quorum sensing, and sporulation functions. Enriched transport proteins included general L-amino acid transport system substrate binding protein (TS-SBP) from ZC4RG25 sp002919265, branched-chain amino acid TS-SBP from ZC4RG17 sp017577575, putative spermidine/putrescine TS-SBP from the family Geminicoccaceae, glutathione TS-SBP from the genus *Glycomyces*, and ferrous iron transport protein B from the genus JADGHX01. Sporulation-related proteins included SASP A from the genus *Planifilum*, the family RSA5, and an unclassified taxon; SASP D from the family ZCTH02-B6; and spore cortex protein from *Geobacillus thermodenitrificans*. Additional enriched proteins included basic membrane protein A from the genus ZC4RG17, trimeric autotransporter adhesin from the genus QMVU01, iron complex outer membrane receptor protein from the genera JACCXG01 and JAJFFL01 and the order Thermaerobacterales, and OmpA-OmpF porin and OOP family proteins from the genera *Abyssalbus*, *Dongia*, *Gelidibacter*, and SZUA-265, the order J063, and an unclassified species. OmpA-OmpF porin production was also elevated in PSB communities under AMF and AMF+PSB treatments across both developmental stages, suggesting that AMF inoculation enhanced porin expression in native or introduced PSB communities (Supplementary Figure 4).

By the fruiting stage, most proteins enriched during flowering remained elevated under co-inoculation, including branched-chain amino acid TS-SBP from ZC4RG17 sp017577575, trimeric autotransporter adhesin from the genus QMVU01, glutamyl endopeptidase from the family RSA9 and an unclassified taxon, basic membrane protein A and related proteins from the genus ZC4RG17, spore coat protein JC from an unclassified taxon, and SASPs A and D from families RSA5 and ZCTH02-B6 (Figure 5), indicating that the co-inoculation effect on rhizosphere gene expression persisted across tomato developmental stages. However, several proteins showed significantly lower abundance during fruiting relative to flowering, including branched-chain amino acid TS-SBP from ZC4RG17 sp017577575 and the genus ZC4RG17, general L-amino acid TS-SBP from ZC4RG25 sp002919265, SASP D from families RSA5, JAAYXC01, and ZCTH02-B6, and SASP A from *Planifilum* (Figure 5D; Supplementary Table 8), reflecting stage-dependent attenuation of specific nutrient transport and sporulation responses.

## Discussion

### Co□inoculation Reshapes the Rhizosphere and Enriches Functionally Diverse Beneficial Taxa

This study addressed the largely unexplored question of how AMF, PSB, and their co-inoculation influence urban green-waste compost microbiome structure and function by integrating metagenomics with metaproteomics across two tomato developmental stages, demonstrating that microbial consortia differentially reshape rhizosphere composition and function in ways that vary both by inoculant treatment and plant developmental stage. Firstly, we observed a significant shift in community structure between the flowering and fruiting stages (Figure 3A), indicating that plant developmental transitions reshaped the rhizomicrobiome. The flowering-to-fruiting transition has been identified as the period of strongest community restructuring in tomato rhizosphere studies, driven by shifts in root-adhering soil from macro– to micro-aggregates that reshape microbial microhabitats^14^. AMF and PSB co-inoculation enriched several beneficial taxa, including Flavobacteriaceae, known for alkaline phosphatase activity and organic P mineralization^15,16^ and Actinobacteriota genera (*Promicromonospora*, *Thermomonospora*, *Thermopolyspora*), which produce antibiotics, siderophores, and hydrolytic enzymes that suppress disease and promote plant vigor^17^. Thermophilic Firmicutes *Geobacillus* and *Planifilum*, linked to antimicrobial volatile organic compound production^18,19^ and S oxidation and bioavailability^20^, were also enriched, consistent with elevated S accumulation in tomato shoots and fruits (Figure 2B). (Figure 2B). Enrichment of the archaeal ammonia oxidizer *Nitrososphaera* further suggests stimulation of nitrification activity in the rhizosphere, potentially expanding nitrate-based N availability and N source diversity for plant uptake, while elevating the risk of nitrate loss due to limited anion retention. Metaproteomic profiles were largely concordant with amplicon sequencing results, with proteins detected from the same taxa enriched under co-inoculation, including streptogrisin D (*Thermomonospora* spp.), glutathione-independent formaldehyde dehydrogenase (*Thermopolyspora* spp.), sporulation-related proteins (*Geobacillus thermodenitrificans*), glyceraldehyde-3-phosphate dehydrogenase and manganese catalase (*Planifilum* spp.), and bacterioferritin (*Nitrososphaera*) (Supplementary Table 6), providing cross-method validation of community composition shifts under co-inoculation (Figure 2B). This enrichment was stronger under co-inoculation than single-inoculation treatments but diminished by fruiting (Figure 3A), consistent with studies in Arabidopsis and Bambara groundnut showing that flowering-stage plants most actively recruit synergistic taxa to meet peak metabolic demands^24,25^. Together, these findings highlight that co-inoculation exerted its strongest influence on rhizomicrobiome composition during flowering, with effects attenuating by fruiting.

### Metaproteomic Insights into Rhizosphere Function under AMF and PSB Co-Inoculation

Across the flowering and fruiting stages, AMF and PSB co-inoculation led to enrichment of microbial transporter proteins mediating uptake of branched-chain amino acids, L-amino acids, glutathione, iron, and sugars, relative to controls (Figure 5B). These multicomponent transporter systems enable bacteria to scavenge nutrients with high affinity and specificity, and can be upregulated under nutrient-limited conditions^26^. Thus, their increased abundance suggests that co-inoculation restructured the rhizosphere in ways that attracted and stimulated microbial taxa capable of responding to new nutrient and signaling cues. Notably, elevated transporter levels reflected not only shifts in microbial community composition but also increased per-cell expression; the branched-chain amino acid SBP from ZC4RG17 sp017577575 and the L-amino acid SBP from ZC4RG25 sp002919265 were substantially more abundant under co-inoculation despite nearly identical MAG abundances across treatments (Figure 5C; Supplementary Table 7), indicating that some taxa increased abundances of specific proteins rather than expanding population size. These patterns in the co-inoculation condition may be the result of a chemically heterogeneous rhizosphere through AMF-driven shifts in root exudate profiles^27,28^, AMF hyphal exudates modulating PSB activity^29,30^, and bacteria-induced changes in root exudate composition^31,32^. Consistent with this, tomato root exudates contain branched-chain amino acids such as valine, leucine, and isoleucine shown to induce catabolism-related gene expression in growth-promoting rhizosphere bacteria^33^, further supporting the view that co-inoculation selectively triggers transporter upregulation in specific microbial groups. This amino acid signaling role is not unique to tomato: in *Arabidopsis*, phenolics and amino acids rather than sugars were the dominant exudate class shaping rhizosphere microbial community composition and functional gene expression across plant development^24^.

Another notable feature of the AMF and PSB co inoculation metaproteome was the elevated production of QS-related proteins (Figure 4C). These included QS proteins such as glutamyl endopeptidase and a putative spermidine/putrescine TS SBP, along with surface interaction proteins such as basic membrane protein A, OmpA-OmpF porins, and trimeric autotransporter adhesins (Figure 5C). Basic membrane protein A and the OmpA-OmpF porins function as surface exposed proteins that mediate cell adhesion, invasion, and evasion of host immune defenses^34–36^, whereas trimeric autotransporter adhesins facilitate bacterial attachment to abiotic surfaces (including plant roots), autoagglutination, and biofilm formation^37^, collectively supporting cell attachment and cell to cell communication. In fact, OmpA-like proteins such as OprF in *Pseudomonas* further function as QS molecule sensors^38^, directly linking surface interaction to QS regulation. This QS-biofilm axis is functionally significant because QS-coordinated biofilms on AMF hyphae have been shown to show enhanced metabolic activities, including elevated indole-3-acetic acid (IAA) production from biofilm-associated bacteria^39^, which promotes lateral root and root hair development and may partly explain the root growth enhancement observed under co-inoculation (Figure 2A). The hyphosphere, a metabolic hotspot enriched in hyphal exudates including formate, acetate, glucose, and amino acids^5,29,30,40^, is the likely site where AMF recruited the taxa responsible for producing these proteins, including QS-related protein-producing members of the order J063, families RSA9 and Geminicoccaceae, and genera ZC4RG17, *Gelidibacter*, SZUA-265, and QMVU01 (Figure 5C). Enrichment of OmpA and OmpF production by *SZUA-265* was stronger during flowering than fruiting (Figure 5D; Supplementary Table 8), consistent with flowering-stage plants most actively recruiting synergistic taxa^24,25^ and suggesting that QS coordination peaks when rhizosphere biochemical complexity is greatest. While physical localization of these taxa within the hyphosphere was not directly verified in this study, elevated OmpA-OmpF porin production by PSB communities under AMF and AMF+PSB treatments across both stages suggests their direct participation in hyphosphere-associated interactions (Supplementary Figure 4).

### Metaproteomic Insights into Developmental Stage-Specific Shifts in Microbial Competitive Pressure and Sporulation Dynamics

Plants strategically shift their root exudate profiles across developmental stages to recruit microbial communities that match their changing nutrient demands^24,41^. In Arabidopsis, early developmental stages are characterized by sugar-dominated exudation supporting broad microbial diversity, while later stages (bolting and flowering) shift toward phenolic and amino acid-enriched profiles that more effectively recruit specific beneficial taxa including N-fixers and PGPRs to meet escalating plant nutrient demands^24^. The ecological rationale for why flowering plants, which have the greatest nutrient demands, simultaneously exude nitrogen-containing compounds into the rhizosphere remains an underexplored research question^42^. Amino acids function as selective chemoattractants for beneficial rhizosphere microorganisms, and other researchers have speculated that their elevated exudation during flowering, when plant energy demand is greatest, could be seen as a strategic recruitment investment that stimulates microbial nutrient mineralization and solubilization, ultimately returning greater nutrient availability to the plant^24,42^. Tomato appears to follow similar developmental exudation patterns, as evidenced by higher production of general L-amino acid TS-SBP by ZC4RG25 sp002919265 and branched-chain amino acid TS-SBP by ZC4RG17 sp017577575 during flowering than fruiting (Figure 5D; Supplementary Table 8), suggesting that plant-derived amino acids may serve as selective recruitment signals for specific beneficial microbial taxa during the high-demand flowering stage. Concurrently, we documented increased sporulation-related protein expression, including SASPs A and D, spore cortex proteins, and spore coat protein JC in co-inoculated treatments across both developmental stages (Figure 5C), though these proteins declined during fruiting (Figure 5D; Supplementary Table 8). This reduction aligns with previously described decreased rhizosphere resource competition as xylem nutrient transport diminishes as tomatoes transition from flowering to fruiting^43^. Taxa expressing sporulation proteins, including families ZCTH02-B6 and RSA5, genus *Planifilum*, and *Geobacillus thermodenitrificans* likely faced competitive stress or resource limitation from other microorganisms and plant processes (Figure 5C), since sporulation typically responds to nutrient-limited, competitively stressful conditions^44^. The case of Planifilum illustrates this complexity: despite population expansion during vegetative growth (Fig. 3C), flowering initiated its sporulation pathways with further SASP A decline at fruiting, demonstrating that taxonomic abundance data alone cannot fully capture real-time microbial functional activity (Figures 5C, 5D)^10,11^.

Overall, these results support AMF and PSB co-inoculation as an effective biological amendment strategy for enhancing rhizosphere microbiome functional activity and nutrient cycling in compost-based Technosols, with direct relevance to sustainable urban agriculture. Our study provides a foundation for future metaproteomic investigations capturing temporal shifts in rhizosphere functional protein dynamics and plant physiological responses, which will deepen mechanistic understanding of how microbial inoculants modulate host-microbiome interactions across crop development.

## Materials and Methods

### Growth Chamber Experiment Design and Plant Management

We established a growth chamber experiment from January to March 2022 at the controlled environment facilities of Cornell University in Ithaca, NY, USA (42°26′49.2″ N, 76°28′37.2″ W). To begin, Brandywine heirloom tomato seeds (Park Seed, USA) were surface-sterilized by immersion in 70% ethanol for 1 minute, followed by treatment with 0.5% sodium hypochlorite for 3 minutes. Each sterilization step was followed by three rinses with sterile distilled deionized (DDI) water to remove residual agents. The seeds were then soaked overnight in a 0.5% solution of Captan 50W fungicide (Hi-Yield, Texas, USA) to eliminate seedborne pathogens, and rinsed thoroughly again with sterile DDI water. Germination was carried out under sterile conditions on autoclaved 8 mm Whatman filter paper discs moistened with sterile DDI water and placed in sterile 9 cm Petri dishes. The dishes were sealed with parafilm and incubated under a 16-hour dark/20°C and 8-hour light/25°C regime for 7 days, achieving a germination rate of 92%. We also prepared a blended compost using compost from the New York City Department of Sanitation (DSNY) compost facility in Staten Island, New York and substrate from the New York City Clean Soil Bank (CSB) at a ratio of 70% compost to 30% CSB sediments. The DSNY composts are derived from leaves, brush, grass clippings, Christmas trees, wood chips, tree stumps, and logs placed in industrial-scale windrows for aerobic, thermophilic composting at temperatures ranging from 50-70°C prior to curing for further organic matter breakdown. The CSB soils are reclaimed excavated sediments from construction sites in New York City for sustainable and safe reuse in urban greening and agriculture projects^45,46^. Following germination, seedlings showing clear epicotyl emergence were transplanted into small pots filled with the prepared compost. These transplants were grown under the same environmental conditions for an additional week to promote acclimation to the potting substrate.

Tomato seedlings selected for compatibility with compost environments were used to establish nine biological replicates per treatment condition, ensuring consistent seedling vigor across experimental groups. The seedlings were two weeks old at the start of the experiment. For AMF inoculation, we used a commercial product, DYNOMYCO Premium Mycorrhizal Inoculant (Mazor, Israel) containing 700 propagules of *Glomus intraradices* and 200 propagules of *Funneliformis mosseae* per gram. The PSB inoculant, Mammoth P (Mammoth, USA), comprised four PSB species: *Enterobacter cloacae*, *Citrobacter freundii*, *Pseudomonas putida*, and *Comamonas testosterone*. The experiment involved 72 tomato plants, evenly split into two groups. One group of 36 plants was grown for an additional four weeks post-germination to reach the flowering stage (six-week-old tomatoes), while the other 36 were cultivated for eight additional weeks to reach the fruiting stage (ten-week-old tomatoes). Each timepoint group was subdivided into four microbial treatments: a control group receiving no AMF or PSB (controls) a PSB group inoculated with PSB only (PSB tomatoes); an AMF group inoculated with AMF only (AMF tomatoes); and a co-inoculated group receiving both AMF and PSB, herein referred to as the AMF+PSB tomato. All 72 tomato plants were cultivated in the prepared compost mixture within a controlled growth chamber, maintained under the same environmental conditions as those used for seed germination. A randomized complete block design was used in the growth chamber to reduce positional environmental bias across treatment groups.

Due to substantial vegetative growth, the tomato plants were transplanted into larger pots following the harvest of the six-week-old plants at the flowering stage. These transplanted individuals were maintained under identical controlled conditions for an additional four weeks to progress into the fruiting stage. Each plant was irrigated exclusively with sterile DDI water to prevent unintended microbial contamination and to maintain the effectiveness of the AMF and PSB inoculations, and it received 50 mL of half strength Hoagland solution twice weekly before flowering, which was increased to 100 mL per application once flowering commenced. Inoculations were performed following the manufacturers’ instructions. AMF was applied initially at 10 g L ¹ of soil volume, and reduced to 5 g L ¹ during later transplanting to adjust for increased pot size. Mammoth P was applied following the manufacturer’s protocol, diluted at a concentration of 0.16 mL per liter and delivered with 50 mL of sterile DDI water. This PSB treatment was administered twice weekly to the PSB and AMF+PSB groups. To ensure consistency in irrigation and handling, the control and AMF groups received an equal volume (50 mL) of sterile DDI water without PSB inoculants.

### Sample Harvest and On-Site Measurements

Tomato samples were harvested at two developmental stages: the flowering phase (six weeks post-germination) and the fruiting stage (ten weeks post-germination). At the flowering stage, tomato roots were sampled to assess AMF colonization, and rhizosphere soil was collected from 36 plants following careful uprooting and removal of bulk soil. At the fruiting stage, an additional set of 36 tomato plants was sampled, from which shoots, roots, and rhizosphere soils were collected using the same procedure. At the fruiting stage, plant biometric measurements were conducted on-site to assess shoot height and maximum root length, serving as key indicators of developmental status during reproductive growth. Plant samples including shoots, roots, and fruits were stored and transported at 4°C in an ice-filled cooler to maintain sample integrity until arrival at the laboratory. Soil samples intended for genomic and metaproteomic analyses, as well as acid phosphatase activity assays, were flash-frozen in liquid nitrogen immediately upon collection and transported to the laboratory on dry ice to minimize protein degradation and maintain microbial functional integrity. Samples were stored at –80 °C until further processing. For nutrient analysis, unripe green fruits were harvested from fruiting-stage plants to assess elemental composition.

### Nutrient Analyses Using ICP-OES

Entire shoots and fruits were oven-dried at 50 °C and finely ground for dry weight determination and subsequent elemental analysis via inductively coupled plasma-optical emission spectrometry (ICP-OES). For fruit nutrient analysis, samples were selected based on size (4-5 cm in diameter) and developmental consistency, standardized to the Breaker stage according to the Dutch Kleurstadia Tomato Color Scale, which marks the onset of physiological ripening when fruits remain fully green without visible yellowing or blush. One fruit per biological replicate was analyzed, yielding 36 representative samples at the fruiting stage (ten-week-old tomatoes). All nutrient analyses were conducted at the Federal Nutrition Laboratory, USDA-ARS Robert W. Holley Center for Agriculture & Health, Ithaca, NY, USA (42°26′52.0″ N, 76°28′03.4″ W).

### AMF Colonization Rate Evaluation

To assess AMF colonization in tomato roots, we employed the ink-vinegar staining technique^47^, following the protocol described by Son et al.^48^. A total of 36 root samples from six week old tomato plants and another 36 from ten week old plants at the fruiting stage were rinsed with distilled water and cleared in 10% potassium hydroxide (KOH) at 80 °C for 30 minutes to remove cellular contents. Subsequently, roots were acidified in 1% hydrochloric acid (HCl) at room temperature for 30 minutes and rinsed again with distilled water. Staining was carried out by boiling the roots in a 5% ink-vinegar solution, prepared using white household vinegar (5% acetic acid), at 80 °C for 30 minutes. After destaining with distilled water, samples were preserved in 50% glycerol and stored at 4 °C until microscopy imaging. For microscopy, 25 root segments per sample were trimmed to 1 cm lengths and mounted on slides with 50% glycerol. Observations were conducted along four cross-sectional transects per slide. The presence of key AMF structures, namely intraradical hyphae, arbuscules, and vesicles, was assessed using the McGonigle et al.^49^ quantification method. Colonization was expressed as the percentage of segments containing AMF structures out of 100 observations per sample.

### GRSP extraction and quantification

GRSP content was quantified following the procedure of Son et al.^48^, using spectrophotometric measurements to determine total glomalin related soil protein (T GRSP). For extraction, 1 g of ground soil stored at 4 °C was repeatedly treated with 8 ml of 50 mM citric acid (pH 8) and autoclaved at 121 °C for 1 h. After each autoclaving cycle, the supernatant was collected by centrifugation at 3,000 rpm for 10 min. Extractions were continued until the solution acquired a yellowish coloration, indicating complete GRSP release. Supernatants from all cycles were pooled and further clarified by centrifugation at 9,000 rpm for 10 min to remove residual soil particles. The combined extracts were quantified using a Bradford assay with bovine serum albumin as the standard. Measurements were performed with the Amplite® Colorimetric Bradford Protein Quantitation Assay Kit (AAA Bioquest, USA) on a Synergy HT Multi Mode Microplate Reader (Agilent Technologies, USA) at 595 nm.

### Soil Acid Phosphatase Assay Protocol

Our protocol was adapted from established methodologies described in previous studies. Detailed procedures and foundational references can be found in German et al.^50^ and Saiya-Cork et al.^51^. To determine soil acid phosphatase activity, an initial pH assessment was conducted on all soil samples, given the enzyme’s high sensitivity to pH variability. Ten grams of soil were mixed with 20 mL of MilliQ water, stirred for 30 minutes, and left to settle at room temperature for one hour to obtain an accurate pH reading. Based on the measured pH, an appropriate buffer was selected: soils with a pH below 7 were treated with a 10X 50 mM sodium acetate buffer (pH ∼5.0, 50 mM), while soils with a neutral to alkaline pH (>7) required a 10X 50 mM sodium bicarbonate buffer (pH 8.0, 50 mM). Our compost-amended soils had relatively high pH values (7.2-7.9), thus necessitating the use of sodium bicarbonate buffer. To generate a standard curve, 1.76 mg of 4-methylumbelliferone (MUB; Sigma-Aldrich M1381, USA) was dissolved in 100 mL of sterile MilliQ water to obtain a 100 µM stock solution, which we handled under reduced light conditions and stored at –20 °C (without refreezing once thawed). Serial dilutions were prepared freshly on the day of the enzyme assay and vortexed briefly. To prepare the fluorogenic substrate, we dissolved 5.12 mg of 4-MUB-phosphate (MUBP; Sigma-Aldrich M8883, USA) in 100 mL of MilliQ water, using moderate heat (< 60 °C) and minimizing light exposure.

Each soil sample (previously stored at –80 °C) was processed by blending 3 g of soil with 150 mL of 1X buffer using a high-speed blender for 30 seconds. A 200 µL aliquot of the resulting slurry was pipetted into a 96-well assay plate, with six technical replicates prepared per sample. To each well, 50 µL of substrate (MUB or MUBP) or buffer was added, and the contents were gently mixed by pipetting ten times; this marked the start of incubation. Plates were incubated for 3 hours at room temperature (22-25 °C) in darkness. Fluorescence was measured immediately after incubation using a pre-warmed microplate fluorometer (Synergy HT Multi-Mode Reader, BioTek, USA) equipped with 365 nm excitation and 450 nm emission filters. For data analysis, we assessed the consistency of technical replicates and applied Dixon’s Q test to remove outliers. A standard curve was generated from MUB, and enzyme activity values were averaged across replicates, excluding any identified outliers. Acid phosphatase activity was calculated in units of nmol g ¹ dry soil h ¹ using the formula:

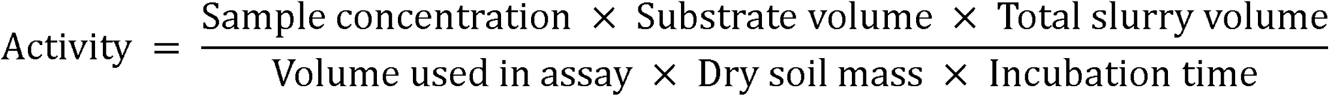

### Statistical Testing of Plant, Soil, and Nutrient Traits

All biometric measurements including plant height, root length, tissue nutrient compositions in shoots and fruits, AMF root colonization percentages, and soil acid phosphatase activity levels were normalized through scaling transformations before conducting statistical analyses. Statistically significant differences in nutrient concentrations between inoculation treatments were identified using pairwise Wilcoxon rank-sum tests executed through the R package stats^52^, with statistical significance established at BH adjusted *p* < 0.05.

### DNA Extraction and Amplicon Sequencing Workflow

Soil DNA was extracted from 0.25 g of frozen soil (stored at −80 °C) using the DNeasy PowerSoil Pro Kit (Qiagen, Carlsbad, CA), following the protocol of Son et al.^48^. The V4 hypervariable region of the 16S rRNA gene was then amplified with universal primers 515F (5’-GTGYCAGCMGCCGCGGTAA-3’) and 806R (5’-GGACTACNVGGGTWTCTAAT-3’), generating amplicons with an average length of 253 base pairs. Each PCR reaction consisted of 10 μl of 2× AccuStart II PCR ToughMix (QuantaBio, Beverly, MA, USA), 1 μl each of the 10 μM forward and reverse primers, 6 μl of DNase-free PCR water (MoBio, Carlsbad, CA, USA), and 2 μl of template DNA. The thermal cycling procedure featured an initial denaturation at 94°C for 2 minutes, followed by 25 cycles of 94°C for 20 seconds, 55°C for 20 seconds, and 72°C for 30 seconds, with a final extension at 72°C for 5 minutes. Library preparation incorporated unique dual-barcode index combinations (Nextera, Illumina) and normalization using clear 96-well plates, following the protocol described by Howard et al.^53^. Finally, the pooled libraries were subjected to paired-end sequencing (2 × 250 bp) with the MiSeq v2 500 bp kit (Illumina, San Diego, CA, USA) at the Cornell Genomics Facility in Ithaca, NY, USA (42°26′47.9″ N, 76°28′42.1″ W).

### Processing of 16S rRNA Gene Amplicon Sequences

The 16S rRNA gene sequencing data were subjected to a standardized bioinformatics pipeline implemented in QIIME2 v.2023.9 (https://qiime2.org)^54^. Following import, amplicon sequence variants (ASVs) were resolved through the DADA2 denoise-paired module, which performs quality filtering, error correction, and merging of paired-end reads^55^. Chimeric sequences were subsequently detected and removed using Uchime, employing an adapted Bayesian algorithm to enhance sensitivity and specificity in chimera identification^56^. The resulting dataset included 522,659 high-quality reads and 14,209 unique amplicon sequence variants (ASVs) associated with the rhizosphere of tomato plants grown in compost-amended substrates. Taxonomic assignments were generated via a Naive Bayes classifier trained on the Greengenes2 2022.10 reference database, using the q2-feature-classifier plugin^57^. Processed ASV tables were then exported to R v4.4.2 for downstream statistical and ecological analysis^52^.

Microbial community composition within rhizosphere samples was characterized using the phyloseq package^52^. Raw ASV counts were transformed to relative abundances, and features with < 0.5% abundance per treatment group were excluded to suppress analytical noise. To improve interpretive robustness, taxa were retained only if present in ≥ 70% of biological replicates within each group, thereby emphasizing ecologically consistent members of the microbiome. Community structure was quantified using Bray-Curtis dissimilarities calculated via the vegdist function in the vegan package^58^ and visualized through PCoA using the ordinate function in phyloseq^59^. To address the compositional constraints intrinsic to amplicon-derived data, ASV matrices underwent centered log-ratio (CLR) transformation using the microbiome package^60^. CLR normalization substantially reduced spurious correlations, thereby enhancing the resolution of community dissimilarity metrics (e.g., Bray-Curtis and Hellinger)^61^. This transformation mitigated compositional distortions inherent to raw relative abundance data, enabling more accurate and interpretable assessments of microbial community structure. Differences in beta diversity across treatment groups were evaluated using permutational multivariate analysis of variance (PERMANOVA), implemented via the adonis2 function in vegan^58^.

### Differential Abundance Analysis of 16S rRNA Gene Amplicon Profiles

Differential abundance analysis was performed on raw ASV count data filtered to retain features present in at least 80% of biological replicates within each treatment group. Analyses were performed using the ANOVA-like Differential Expression (ALDEx) framework implemented in the ALDEx2 R package^13^, which has been demonstrated to yield high analytical fidelity by minimizing false discovery rates in compositional microbiome data^62,63^. ALDEx2 addresses the compositional nature of sequencing data by applying a CLR transformation to the count matrix, thereby projecting the data into an unconstrained Euclidean space. The pipeline generated 128 Monte Carlo Dirichlet instances per sample, simulating posterior distributions of CLR-transformed ASV counts. Statistical inference between groups was performed using the Wilcoxon rank-sum test, and significance was determined based on BH adjusted *p* values with a false discovery threshold of 0.05.

### Soil Protein Extraction and Peptide Preparation

We extracted proteins and digested them into peptides for mass spectrometry analysis using a combined precipitation and filter aided sample preparation (FASP) method^64–66^. Proteins were extracted from 200 mg of frozen soil (stored at –80□°C) using bead-beating-assisted lysis in 2□mL Lysing Matrix E tubes (MP Biomedicals, cat#: 6914050, USA) supplemented with 1.2□mL SDT lysis buffer (4% w/v SDS, 100□mM Tris-HCl, pH□7.6, 0.1□M DTT). Mechanical lysis was performed via bead beating at 6 m/s for three 45-second cycles with 30-second dwell intervals between each cycle. Lysates were heated at 95□°C for 10□min to enhance protein solubilization, followed by centrifugation at 17,000□×□g for 10□min. Supernatants were transferred to fresh tubes and further clarified by sequential centrifugation at 17,000□×□g and 18,000□×□g for 10□min each. Proteins were precipitated at 25% (w/v) trichloroacetic acid (TCA) and stored overnight at –80□°C. Following thawing, samples were centrifuged at 21,000□×□g for 15□min at 4□°C. Resulting pellets were washed with 1 mL ice-cold acetone, vortexed, and centrifuged again (21,000□×□g, 15□min, 4□°C). Pellets were air-dried in inverted tubes, and then were resuspended in 60 μL SDT lysis buffer. Prior to loading the resuspended pellets, 10kDa filter units (Microcon® Centrifugal Filters, Millipore, US) were washed with 50□μL of 0.1% formic acid (FA) and centrifuged at 14,000□×□g for 5□min, followed by sequential washes with 50 μL of 50□mM ammonium bicarbonate (ABC) and 50□μL of NaCl. The resuspended pellets were loaded onto the filter and mixed with 200□μL UA buffer (8□M urea, 0.1□M Tris-HCl, pH□8.5), and centrifuged at 14,000□×□g for 40□min. Proteins were alkylated with 100□μL of iodoacetamide (IAA; 0.05% in UA buffer), mixed at 600□rpm for 1□min, incubated in darkness for 20□min, and centrifuged (14,000□×□g, 30-40□min). Filters were washed three times with UA buffer (100 μL, centrifuged at 14,000 × g, 15 min). A buffer exchange was performed through three subsequent washes with ABC buffer (100 μL, centrifuged at 14,000 × g, 15 min). Then, proteins were digested by adding 0.5□μg of Pierce™ MS-Grade Trypsin Protease (Thermo Scientific, USA) reconstituted in 40 μL ABC buffer, mixing at 600□rpm for 1□min, and incubating overnight (15-18□h) at 37□°C in a humidified chamber. Resulting peptides were recovered by centrifugation at 14,000□×□g for 20□min and a subsequent wash with 50□μL of 0.5□M NaCl (mixed at 600□rpm for 1□min, centrifuged 20□min). Peptide yield was quantified using the Pierce Micro BCA Protein Assay Kit (Thermo Scientific, USA) following the manufacturer’s guidelines.

### LC-MS/MS Analysis of Soil-Derived Peptides

Liquid chromatography-tandem mass spectrometry (LC-MS/MS) was conducted following the methods previously described by Violette et al.^65^ Peptides (1,000 ng) were loaded onto a 5□mm × 300□μm i.d. Acclaim PepMap100 C18 trap column (Thermo Fisher Scientific, USA) using loading solvent A (2% acetonitrile, 0.05% trifluoroacetic acid) via an UltiMate™ 3000 RSLCnano system (Thermo Fisher Scientific, USA). Peptides were then eluted onto an EASY-Spray C18 analytical column (PepMap RSLC, 2□μm, 75□cm × 75□μm) maintained at 60□°C for separation. Peptides were separated using a 140-minute gradient at a flow rate of 300 nL/min. The gradient proceeded from 5% mobile phase B (0.1% formic acid in water) to 31% mobile phase B (0.1% formic acid in 80% acetonitrile) over 102 minutes, increased to 50% B over the next 18 minutes, and was held at 99% B for a final 20-minute wash. The separation column was attached to an Orbitrap Exploris 480 Mass Spectrometer (ThermoFisher Scientific, USA) via an Easy-Spray source where eluting peptides were ionized by electrospray ionization. Full MS scans were performed to acquire MS^1^ spectra over a 380-1600 m/z scan range at 60,000 resolutions with a maximum injection time of 200 ms and a normalized AGC Target of 300% (3e6). MS^2^ spectra were acquired for the top 15 most abundant ions. Ions were isolated with a maximum injection time of 50 ms and a normalized AGC target of 100% (1e5), then subjected to a normalized collision energy of 27% applied in the HCD cell to generate peptide fragments. MS^2^ scans of peptide fragments were acquired at 15,000 resolutions and ions excluded from additional MS^2^ for 25 s. A column wash method, which oscillated between 5-99% mobile phase B every 3-4 minutes for 30 minutes, was incorporated between sample runs to minimize sample carryover.

### Metagenomic Processing and Sample-Matched Metaproteome Database Construction

To enable *de novo* construction of a metaproteomic search database from environmental microbiomes where microbial population is unknown and not previously sequenced, we assembled a genome-resolved metagenome from shotgun metagenomic reads as the basis for the protein database for metaproteomics^67,68^. The control and AMF and PSB co-inoculation treatment groups were selected for sequencing to capture both the baseline activity of native soil microbial communities and the functional influence of exogenously introduced microbial inoculants. The co□inoculated metagenome already contained the taxa present in the single□inoculation treatments, so sequencing the AMF or PSB tomato groups would not have contributed additional information for constructing the sample□matched protein database. For each developmental stage (flowering and fruiting), we collected three biological replicates from two treatments (control and AMF+PSB), resulting in four total sample groups (flowering□control, flowering□AMF+PSB, fruiting□control, and fruiting□AMF+PSB). All triplicates were subjected to shallow sequencing, and one pooled sample per group was selected for deep sequencing to enhance metagenome coverage and detect low-abundance features. Shallow sequencing facilitated differential taxonomic abundance testing, while deep sequencing provided the coverage necessary for robust genome-centric assembly. For shallow sequencing, genomic DNA was normalized to 25□ng/μL using NanoDrop spectrophotometry (Thermo Fisher Scientific, USA) to ensure consistent input mass. All metagenomic samples were processed at the Genomic Sciences Laboratory (North Carolina State University, Raleigh, NC, USA) (35°47′12.3″□N, 78°40′19.5″□W). Sequencing was performed on three NovaSeq 6000 S4 lanes (Illumina, 150□bp paired-end), with shallow libraries allocated to one lane and deep-sequencing libraries to the remaining two lanes. High□throughput sequencing yielded an average of ∼71□million (shallow) to ∼541□million (deep) paired end reads per sample, with total base counts averaging 10.5-81.7□Gb. All libraries exhibited a consistent read length of 151 bp and an average GC content of ∼64-65%. Prior to sequencing, sample integrity and fragment size distribution were verified using TapeStation High Molecular Weight (HMW) DNA analysis to ensure suitability for downstream metagenomic and metaproteomic applications.

Read quality control was performed with FastQC v.0.12.1^69^, including per-base sequence quality, GC composition, adapter contamination, and duplication metrics. To obtain a preliminary overview of the organisms present, PhyloFlash v.138.1^70^ was used. Adapter trimming and removal of PhiX174 spike-in reads (GenBank CP004084.1) were performed with BBDuk (BBMap v.39.01)^71^ using parameters mink = 6 and minlength = 20. To remove exogenous microbial and host-derived reads, BBSplit (BBMap v.39.01)^71^ was used to filter sequences matching reference genomes from the NCBI GenBank database. This included four PSB species present in the Mammoth P inoculant (Mammoth, US): *Enterobacter cloacae* (GCA_905331265.2), *Citrobacter freundii* (GCA_904859905.1), *Pseudomonas putida* (GCA_000412675.1), and *Comamonas testosteroni* (AAUJ02000001.1). As an assembled genome for *Comamonas testosteroni* was unavailable, shotgun sequencing reads (fragmented) were used to ensure accurate masking and downstream resolution. Similarly, genomes of AMF strains used in the DYNOMYCO Premium Mycorrhizal Inoculant (DYNOMYCO, Israel) were masked, including *Rhizophagus irregularis* (GCA_026210795.1) and *Funneliformis mosseae* (GCA_910592005.1). Because the genome of the Brandywine tomato cultivar has not yet been sequenced, we assembled a surrogate host genome reference from publicly available tomato germplasm encompassing twenty-one wild and cultivated accessions (GCA_025759855.1).

Assembly of filtered reads was conducted using MEGAHIT v.1.2.9^72^ with k-mer sizes of 33, 55, 77, 99, 119, and 141. Shallow and deep sequencing data were co-assembled at the farm level to construct high-resolution metagenomic references per site. Assembly quality was assessed with QUAST v.5.2.0^73^. All raw reads were then mapped to the assembled contigs using BBMap to obtain coverage profiles for binning^71^, followed by reconstruction of metagenome assembled genomes (MAGs) with MetaBAT2 v.2.15^74^. Bins were assessed for completeness and contamination using CheckM v.1.0.1^75^, following established MAG quality criteria^12^. MAGs with ≥ 50% completeness and < 10% contamination were considered medium to high-to-medium quality; all other bins retained as low quality, following the criteria established by Bowers et al.^12^ (Supplementary Table 9). Bins exceeding 10% contamination were excluded, as maintaining contamination levels below this threshold ensures the recovery of cleaner and more taxonomically consistent MAGs. Species-level dereplication was performed using dRep v3.4.2^76^ with a 95% average nucleotide identity (ANI) threshold to group redundant MAGs. Final taxonomic assignments were generated using GTDB-Tk v.2.4.0^77^ against the GTDB r220 release, providing standardized phylogenomic classification of prokaryotic MAGs. To complement this, BAT v5.2.3^78^ using the NCBI database was employed to support taxonomic resolution for prokaryotes MAGs not covered by the GTDB database. A non-redundant metaproteome search database was constructed by integrating protein-coding sequences predicted from MAGs using Prodigal v.2.6.3^79^ and clustered with CD-HIT v.4.8.1^80^ at 95% sequence identity. To further minimize redundancy, we first prioritized proteins from medium□ and high□ quality MAGs and then used cd□ hit□ 2d to identify and add non□ redundant proteins from low□ quality MAGs and unbinned contigs that were missing from the higher□ quality set. The finalized database was merged with reference proteomes from tomato, AMF, and PSB strains, and appended with cRAP contaminants (e.g., keratins, trypsin) from the common Repository of Adventitious Proteins (https://www.thegpm.org/crap/), facilitating downstream spectral annotation.

### Differential Abundance Analysis of MAG Profiles

To assess differential taxonomic abundance, shallow metagenomic datasets from three biological replicates of both the control and AMF+PSB-treated groups per developmental stage were processed using BBsplit (BBMap v.39.01)^71^. Reads were aligned to reference genome collections with a stringent identity threshold of 97%, enabling high-confidence taxonomic assignment. Genome-level mapping statistics were generated via the refstats function, while unmapped reads were retained in separate FASTQ files for downstream analysis. Relative abundances were calculated based on the proportion of uniquely mapped reads. Statistical comparisons of taxon-level abundance profiles were conducted in R v.4.4.2^52^, using the rstatix package^81^. Welch’s two-sample t-tests assuming unequal variances were applied, and significance was determined using BH adjusted *p* values (BH adjusted *p* < 0.05).

### Metaproteomic Analysis and Functional Annotation Workflow

We identified proteins by searching MS/MS spectra against the custom protein sequence database using Proteome Discoverer v2.3 (Thermo Fisher Scientific, USA), following the workflow described in Violette et al.^65^. Peptides were identified using the Sequest HT and percolator nodes with the following parameters: trypsin digestion (full specificity) allowing up to two missed cleavages, precursor mass tolerance of 10 ppm, fragment ion tolerance of 0.1 Da, and permitting up to three dynamic modifications per peptide. Protein inference was implemented via the Protein FDR Percolator node, enforcing dual thresholds of 0.01 and 0.05 to maintain a false discovery rate (FDR) below 5%. Search results from all samples were combined into a multiconsensus report, and only Master proteins identified with medium or high confidence were retained. Protein abundances were quantified using normalized spectral abundance factors (NSAFs), which were multiplied by 100 to obtain relative protein abundances (%). For functional annotation, significant proteins were annotated using multiple annotation pipelines, including Mantis (v.1.5.5)^82^, eggNOG-mapper (v.2.1.12)^83^, Blastkoala2 and GhostKoala^84^, and KofamKoala^85^. Only proteins with consensus KEGG Orthology (KO) assignments were retained to enhance classification reliability. Final pathway enrichment and hierarchical functional annotation (three KEGG pathway levels) were conducted using the R package KEGGREST (v.1.46.0)^86^, allowing for mapping of detected proteins to biological pathways.

### Visualization of Functional Variation Across Agricultural Sites

Quantitative proteomic data were subsequently imported into R v.4.4.2^52^ for statistical analysis. Proteins detected in ≥ 70% of samples within each treatment group were retained for downstream analysis to minimize sampling bias. Abundance values were NSAF and log□ transformed, with zero or infinite values replaced by an imputed minimum value of –1 to maintain continuity in downstream analyses. Differential abundance between treatments was assessed using a Welch’s two-sample t-test implemented via the rstatix package^81^, with *p* values adjusted for multiple testing using the BH procedure. For multivariate analyses, raw abundance counts were filtered using the same ≥ 70% prevalence threshold and normalized with a CLR transformation to account for compositional structure, with zeros replaced by the minimum non zero value × 0.1 prior to transformation. Principal component analysis (PCA) was performed using the prcomp function in the stats package (v.4.4.2)^52^, and resulting ordinations were visualized using ggplot2 (v.3.5.1)^87^. Protein compositional differences among treatment groups were evaluated using pairwise PERMANOVA on Euclidean distance matrices derived from CLR transformed abundance values. Statistical significance was determined at a BH adjusted *p* < 0.05. Volcano plots were generated using the R package ggplot2 (v3.5.1)^87^ to visualize overall differential expression patterns, highlighting genes based on log□ fold change and statistical significance (BH adjusted *p* < 0.1). To further dissect treatment– and stage-specific differences, protein profiles from pairwise comparisons across two developmental stages (flowering vs. fruiting) and four inoculation treatments were visualized as heatmaps using the pheatmap package (v1.0.13)^88^.

## Acknowledgments

The authors gratefully acknowledge Sammi Lin and Yuzhi Wang for their outstanding contributions to tomato cultivation and sampling. We also thank Zachary Butler-Jones for his invaluable assistance with the demanding work of securing subsoil sand, harvesting, and sampling. All LC–MS/MS measurements were made in the Molecular Education, Technology, and Research Innovation Center (METRIC) at North Carolina State University.

## Author contributions

Conceptualization: YS, JKK, MK

Methodology: JKK, EJC, MAP, MK

Investigation: YS, MK

Visualization: YS

Supervision: MK, JKK

Writing—original draft: YS

Writing—review & editing: EJC, MAP, OM, MK, JKK

## Data and materials availability

Metagenomic reads, medium– to high-quality metagenome-assembled genomes, and 16S rRNA amplicon sequences generated in this study are publicly available in the NCBI Sequence Read Archive (SRA) under BioProject accession number PRJNA1443301. Datasets, R scripts, and processed outputs generated in this study have been made publicly available in the following GitHub repository: https://github.com/noel1105/Tomato_Developmental_metaproteome. Proteomic data, raw mass spectrometry files, and databases used are deposited in the ProteomeXchange Consortium via the PRIDE partner repository (REF)^89,90^ under the project accessions PXD077937 [REVIEWER TOKEN: inWQN8AMzfjm (https://www.ebi.ac.uk/pride/login)]).

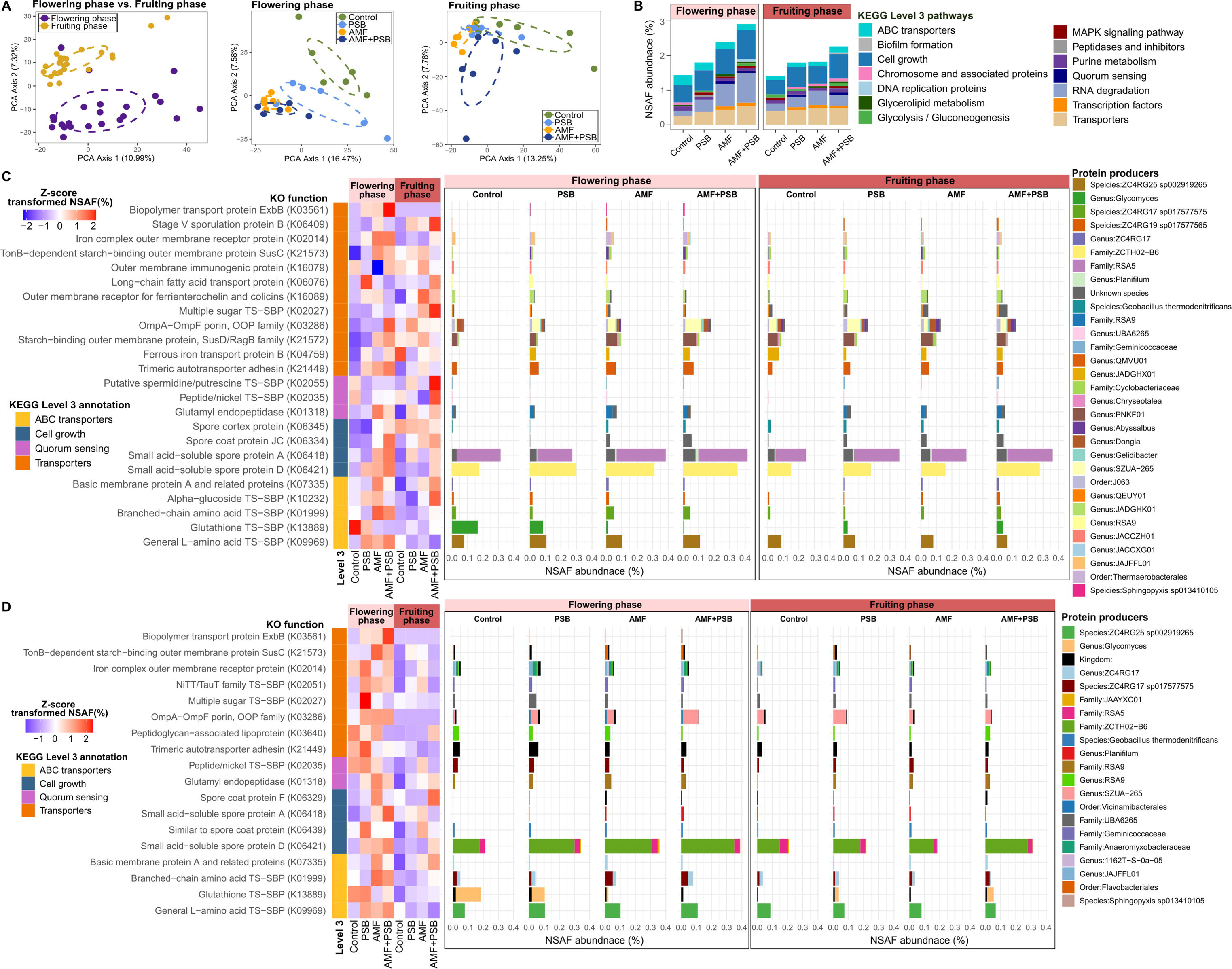

## Reference

1. Schröder, A., Schloter, M., Roccotiello, E., Weisser, W. W. & Schulz, S. Improving ecosystem services of urban soils – how to manage the microbiome of Technosols? Front. Environ. Sci. 12, (2024).

2. Trivedi, P., Mattupalli, C., Eversole, K. & Leach, J. E. Enabling sustainable agriculture through understanding and enhancement of microbiomes. New Phytologist 230, 2129–2147 (2021).

3. Sharma, M. et al. AMF and PSB applications modulated the biochemical and mineral content of the eggplants. Journal of Basic Microbiology 62, 1371–1378 (2022).

4. Ordoñez, Y. M. et al. Bacteria with Phosphate Solubilizing Capacity Alter Mycorrhizal Fungal Growth Both Inside and Outside the Root and in the Presence of Native Microbial Communities. PLOS ONE 11, e0154438 (2016).

5. Duan, S. et al. Cross-kingdom nutrient exchange in the plant–arbuscular mycorrhizal fungus–bacterium continuum. Nat Rev Microbiol 22, 773–790 (2024).

6. Zhang, C. et al. A tripartite bacterial-fungal-plant symbiosis in the mycorrhiza-shaped microbiome drives plant growth and mycorrhization. Microbiome 12, 13 (2024).

7. El Maaloum, S., et al. Effect of Arbuscular Mycorrhizal Fungi and Phosphate-Solubilizing Bacteria Consortia Associated with Phospho-Compost on Phosphorus Solubilization and Growth of Tomato Seedlings (Solanum lycopersicum L.). Communications in Soil Science and Plant Analysis 51, 622–634 (2020).

8. El Maaloum, S., et al. Effect of Arbuscular Mycorrhizal Fungi and Phosphate-Solubilizing Bacteria Consortia Associated with Phospho-Compost on Phosphorus Solubilization and Growth of Tomato Seedlings (Solanum lycopersicum L.). Communications in Soil Science and Plant Analysis 51, 622–634 (2020).

9. Tahiri, A. et al. Assessing the Potential Role of Compost, PGPR, and AMF in Improving Tomato Plant Growth, Yield, Fruit Quality, and Water Stress Tolerance. J Soil Sci Plant Nutr 22, 743–764 (2022).

10. Kleiner, M. Metaproteomics: Much More than Measuring Gene Expression in Microbial Communities. mSystems 4, 10.1128/msystems.00115-19 (2019).

11. Zampieri, E., Chiapello, M., Daghino, S., Bonfante, P. & Mello, A. Soil metaproteomics reveals an inter-kingdom stress response to the presence of black truffles. Sci Rep 6, 25773 (2016).

12. Bowers, R. M. et al. Minimum information about a single amplified genome (MISAG) and a metagenome-assembled genome (MIMAG) of bacteria and archaea. Nat Biotechnol 35, 725–731 (2017).

13. Fernandes, A. D., Macklaim, J. M., Linn, T. G., Reid, G. & Gloor, G. B. ANOVA-Like Differential Expression (ALDEx) Analysis for Mixed Population RNA-Seq. PLOS ONE 8, e67019 (2013).

14. Dong, M. et al. Tomato growth stage modulates bacterial communities across different soil aggregate sizes and disease levels. ISME Communications 3, 104 (2023).

15. Cao, J., Huang, Y. & Wang, C. Rhizosphere interactions between earthworms (Eisenia fetida) and arbuscular mycorrhizal fungus (Funneliformis mosseae) promote utilization efficiency of phytate phosphorus in maize. Applied Soil Ecology 94, 30–39 (2015).

16. Gavriilidou, A. et al. Comparative genomic analysis of Flavobacteriaceae: insights into carbohydrate metabolism, gliding motility and secondary metabolite biosynthesis. BMC Genomics 21, 569 (2020).

17. Barka, E. A., et al. Taxonomy, Physiology, and Natural Products of Actinobacteria. Microbiology and Molecular Biology Reviews: MMBR 80, 1 (2015).

18. McMullan, G. et al. Habitat, applications and genomics of the aerobic, thermophilic genus Geobacillus. Biochemical Society Transactions 32, 214–217 (2004).

19. Han, S.-I., Lee, J.-C., Lee, H.-J. & Whang, K.-S. Planifilum composti sp. nov., a thermophile isolated from compost. International Journal of Systematic and Evolutionary Microbiology 63, 4557–4561 (2013).

20. Ren, Y., Strobel, G., Sears, J. & Park, M. Geobacillus sp., a Thermophilic Soil Bacterium Producing Volatile Antibiotics. Microb Ecol 60, 130–136 (2010).

21. Huang, L. et al. Ammonia-oxidizing archaea are integral to nitrogen cycling in a highly fertile agricultural soil. ISME COMMUN. 1, 19 (2021).

22. Heyman, H., Bassuk, N., Bonhotal, J. & Walter, T. Compost Quality Recommendations for Remediating Urban Soils. International Journal of Environmental Research and Public Health 16, 3191 (2019).

23. Wright, A. L., Provin, T. L., Hons, F. M., Zuberer, D. A. & White, R. H. Compost impacts on dissolved organic carbon and available nitrogen and phosphorus in turfgrass soil. Waste Management 28, 1057–1063 (2008).

24. Chaparro, J. M., Badri, D. V. & Vivanco, J. M. Rhizosphere microbiome assemblage is affected by plant development. ISME J 8, 790–803 (2014).

25. Ajilogba, C. F., Olanrewaju, O. S. & Babalola, O. O. Plant Growth Stage Drives the Temporal and Spatial Dynamics of the Bacterial Microbiome in the Rhizosphere of Vigna subterranea. Front. Microbiol. 13, (2022).

26. Davies, J. S. et al. Selective Nutrient Transport in Bacteria: Multicomponent Transporter Systems Reign Supreme. Front. Mol. Biosci. 8, (2021).

27. Ma, J. et al. AMF colonization affects allelopathic effects of Zea mays L. root exudates and community structure of rhizosphere bacteria. Front. Plant Sci. 13, (2022).

28. Hage-Ahmed, K., Moyses, A., Voglgruber, A., Hadacek, F. & Steinkellner, S. Alterations in Root Exudation of Intercropped Tomato Mediated by the Arbuscular Mycorrhizal Fungus Glomus mosseae and the Soilborne Pathogen Fusarium oxysporum f.sp. lycopersici. Journal of Phytopathology 161, 763–773 (2013).

29. Kalamulla, R. & Yapa, N. Co-inoculation of AMF and Other Microbial Biofertilizers for Better Nutrient Acquisition from the Soil System. in Arbuscular Mycorrhizal Fungi in Sustainable Agriculture: Nutrient and Crop Management (eds Parihar, M., Rakshit, A., Adholeya, A. & Chen, Y.) 99–111 (Springer Nature, Singapore, 2024). doi:10.1007/978-981-97-0300-5_4.

30. Duan, S., Zhang, L. & Declerck, S. Early-stage reciprocal cooperation between the arbuscular mycorrhizal fungus Rhizophagus irregularis and the phosphate-solubilizing bacterium Rahnella aquatilis is dependent on external phosphorus availability. Commun Biol 8, 1075 (2025).

31. Zhang, H. et al. Certain Tomato Root Exudates Induced by Pseudomonas stutzeri NRCB010 Enhance Its Rhizosphere Colonization Capability. Metabolites 13, 664 (2023).

32. Krzyżanowska, D. M. et al. Host-adaptive traits in the plant-colonizing Pseudomonas donghuensis P482 revealed by transcriptomic responses to exudates of tomato and maize. Sci Rep 13, 9445 (2023).

33. Krzyżanowska, D. M. et al. Host-adaptive traits in the plant-colonizing Pseudomonas donghuensis P482 revealed by transcriptomic responses to exudates of tomato and maize. Scientific Reports 13, 9445 (2023).

34. Confer, A. W. & Ayalew, S. The OmpA family of proteins: Roles in bacterial pathogenesis and immunity. Veterinary Microbiology 163, 207–222 (2013).

35. Choi, U. & Lee, C.-R. Distinct Roles of Outer Membrane Porins in Antibiotic Resistance and Membrane Integrity in Escherichia coli. Front. Microbiol. 10, (2019).

36. Bryksin, A. V., Tomova, A., Godfrey, H. P. & Cabello, F. C. BmpA is a surface-exposed outer-membrane protein of Borrelia burgdorferi. FEMS Microbiol Lett 309, 77–83 (2010).

37. Okaro, U., Green, R., Mohapatra, S. & Anderson, B. The trimeric autotransporter adhesin BadA is required for in vitro biofilm formation by Bartonella henselae. npj Biofilms Microbiomes 5, 10 (2019).

38. Krishnan, S. & Prasadarao, N. V. Outer membrane protein A and OprF: versatile roles in Gram-negative bacterial infections. The FEBS Journal 279, 919–931 (2012).

39. Sharma, S., Compant, S., Ballhausen, M.-B., Ruppel, S. & Franken, P. The interaction between Rhizoglomus irregulare and hyphae attached phosphate solubilizing bacteria increases plant biomass of Solanum lycopersicum. Microbiological Research 240, 126556 (2020).

40. Pandit, A., Adholeya, A., Cahill, D., Brau, L. & Kochar, M. Microbial biofilms in nature: unlocking their potential for agricultural applications. Journal of Applied Microbiology 129, 199–211 (2020).

41. Zhao, M. et al. Root exudates drive soil-microbe-nutrient feedbacks in response to plant growth. Plant, Cell & Environment 44, 613–628 (2021).

42. Moe, L. A. Amino acids in the rhizosphere: From plants to microbes. American Journal of Botany 100, 1692–1705 (2013).

43. Bodale, I. et al. Evaluation of the Nutrients Uptake by Tomato Plants in Different Phenological Stages Using an Electrical Conductivity Technique. Agriculture 11, 292 (2021).

44. Gray, D. A. et al. Extreme slow growth as alternative strategy to survive deep starvation in bacteria. Nat Commun 10, 890 (2019).

45. Garcia, J. et al. Plant growth and microbial responses from urban agriculture soils amended with excavated local sediments and municipal composts. Journal of Urban Ecology 9, juad016 (2023).

46. Walsh, D. et al. Sediment exchange to mitigate pollutant exposure in urban soil. Journal of Environmental Management 214, 354–361 (2018).

47. Vierheilig, H., Coughlan, A. P., Wyss, U. & Piché, Y. Ink and Vinegar, a Simple Staining Technique for Arbuscular-Mycorrhizal Fungi. Appl. Environ. Microbiol. 64, 5004 (1998).

48. Son, Y. et al. Synergistic enhancement of Sorghum bicolor nutrient uptake and growth by microbiomes in enhanced biological phosphorus removal system and arbuscular mycorrhizal fungi. Environmental Microbiome 20, 155 (2025).

49. McGONIGLE, T. P., Miller, M. H., Evans, D. G., Fairchild, G. L. & Swan, J. A. A new method which gives an objective measure of colonization of roots by vesicular—arbuscular mycorrhizal fungi. New Phytologist 115, 495–501 (1990).

50. German, D. P. et al. Optimization of hydrolytic and oxidative enzyme methods for ecosystem studies. Soil Biology and Biochemistry 43, 1387–1397 (2011).

51. Saiya-Cork, K. R., Sinsabaugh, R. L. & Zak, D. R. The effects of long term nitrogen deposition on extracellular enzyme activity in an Acer saccharum forest soil. Soil Biology and Biochemistry 34, 1309–1315 (2002).

52. R Core Team. R: A Language and Environment for Statistical Computing. https://www.r-project.org/ (2024).

53. Howard, M. M., Bell, T. H. & Kao-Kniffin, J. Soil microbiome transfer method affects microbiome composition, including dominant microorganisms, in a novel environment. FEMS Microbiol Lett 364, fnx092 (2017).

54. Bolyen, E. et al. Reproducible, interactive, scalable and extensible microbiome data science using QIIME 2. Nat Biotechnol 37, 852–857 (2019).

55. Callahan, B. J. et al. DADA2: High-resolution sample inference from Illumina amplicon data. Nat Methods 13, 581–583 (2016).

56. Edgar, R. C., Haas, B. J., Clemente, J. C., Quince, C. & Knight, R. UCHIME improves sensitivity and speed of chimera detection. Bioinformatics 27, 2194–2200 (2011).

57. McDonald, D. et al. Greengenes2 enables a shared data universe for microbiome studies. 2022.12.19.520774 Preprint at 10.1101/2022.12.19.520774 (2022).

58. Dixon, P. VEGAN, a package of R functions for community ecology. J Veg Sci 14, 927–930 (2003).

59. McMurdie, P. J. & Holmes, S. phyloseq: An R Package for Reproducible Interactive Analysis and Graphics of Microbiome Census Data. PLOS ONE 8, e61217 (2013).

60. Leo, L. & Sudarshan, S. microbiome R package. Bioconductor 10.18129/B9.bioc.microbiome (2017).

61. Gloor, G. B., Macklaim, J. M., Pawlowsky-Glahn, V. & Egozcue, J. J. Microbiome Datasets Are Compositional: And This Is Not Optional. Front. Microbiol. 8, (2017).

62. McMurdie, P. J. & Holmes, S. Waste Not, Want Not: Why Rarefying Microbiome Data Is Inadmissible. PLOS Computational Biology 10, e1003531 (2014).

63. Nearing, J. T. et al. Microbiome differential abundance methods produce different results across 38 datasets. Nat Commun 13, 342 (2022).

64. Qian, C. & Hettich, R. L. Optimized Extraction Method To Remove Humic Acid Interferences from Soil Samples Prior to Microbial Proteome Measurements. J Proteome Res 16, 2537–2546 (2017).

65. Violette, M. J. et al. Meta-omics reveals role of photosynthesis in microbially induced carbonate precipitation at a CO2-rich geyser. ISME Commun 4, ycae139 (2024).

66. Wiśniewski, J. R., Zougman, A., Nagaraj, N. & Mann, M. Universal sample preparation method for proteome analysis. Nat Methods 6, 359–362 (2009).

67. Bartlett, A., Blakeley-Ruiz, J. A., Richie, T., Theriot, C. M. & Kleiner, M. Large Quantities of Bacterial DNA and Protein in Common Dietary Protein Source Used in Microbiome Studies. PROTEOMICS n/a, e202400149 (2025).

68. Blakeley-Ruiz, J. A. & Kleiner, M. Considerations for constructing a protein sequence database for metaproteomics. Computational and Structural Biotechnology Journal 20, 937–952 (2022).

69. Andrews, S. FastQC: A Quality Control Tool for High Throughput Sequence Data [Online]. http://www.bioinformatics.babraham.ac.uk/projects/fastqc/ (2010).

70. Gruber-Vodicka, H. R., Seah, B. K. B. & Pruesse, E. phyloFlash: Rapid Small-Subunit rRNA Profiling and Targeted Assembly from Metagenomes. mSystems 5, 10.1128/msystems.00920-20 (2020).

71. Bushnell, B. BBMap short read aligner, and other bioinformatic tools. (2016).

72. Li, D., Liu, C.-M., Luo, R., Sadakane, K. & Lam, T.-W. MEGAHIT: an ultra-fast single-node solution for large and complex metagenomics assembly via succinct de Bruijn graph. Bioinformatics 31, 1674–1676 (2015).

73. Gurevich, A., Saveliev, V., Vyahhi, N. & Tesler, G. QUAST: quality assessment tool for genome assemblies. Bioinformatics 29, 1072–1075 (2013).

74. Kang, D. D. et al. MetaBAT 2: an adaptive binning algorithm for robust and efficient genome reconstruction from metagenome assemblies. PeerJ 7, e7359 (2019).

75. Parks, D. H., Imelfort, M., Skennerton, C. T., Hugenholtz, P. & Tyson, G. W. CheckM: assessing the quality of microbial genomes recovered from isolates, single cells, and metagenomes. Genome Res 25, 1043–1055 (2015).

76. Olm, M. R., Brown, C. T., Brooks, B. & Banfield, J. F. dRep: a tool for fast and accurate genomic comparisons that enables improved genome recovery from metagenomes through de-replication. The ISME Journal 11, 2864–2868 (2017).

77. Chaumeil, P.-A., Mussig, A. J., Hugenholtz, P. & Parks, D. H. GTDB-Tk v2: memory friendly classification with the genome taxonomy database. Bioinformatics 38, 5315–5316 (2022).

78. von Meijenfeldt, F. A. B., Arkhipova, K., Cambuy, D. D., Coutinho, F. H. & Dutilh, B. E. Robust taxonomic classification of uncharted microbial sequences and bins with CAT and BAT. Genome Biology 20, 217 (2019).

79. Hyatt, D. et al. Prodigal: prokaryotic gene recognition and translation initiation site identification. BMC Bioinformatics 11, 119 (2010).

80. Li, W. & Godzik, A. Cd-hit: a fast program for clustering and comparing large sets of protein or nucleotide sequences. Bioinformatics 22, 1658–1659 (2006).

81. Kassambara, A. rstatix: Pipe-Friendly Framework for Basic Statistical Tests. https://rpkgs.datanovia.com/rstatix/ (2023).

82. Queirós, P., Delogu, F., Hickl, O., May, P. & Wilmes, P. Mantis: flexible and consensus-driven genome annotation. GigaScience 10, giab042 (2021).

83. Cantalapiedra, C. P., Hernández-Plaza, A., Letunic, I., Bork, P. & Huerta-Cepas, J. eggNOG-mapper v2: Functional Annotation, Orthology Assignments, and Domain Prediction at the Metagenomic Scale. Mol Biol Evol 38, 5825–5829 (2021).

84. Kanehisa, M., Sato, Y. & Morishima, K. BlastKOALA and GhostKOALA: KEGG Tools for Functional Characterization of Genome and Metagenome Sequences. Journal of Molecular Biology 428, 726–731 (2016).

85. Aramaki, T. et al. KofamKOALA: KEGG Ortholog assignment based on profile HMM and adaptive score threshold. Bioinformatics 36, 2251–2252 (2020).

86. Tenenbaum, D. & Maintainer, B. KEGGREST: Client-side REST access to the Kyoto Encyclopedia of Genes and Genomes (KEGG). R package version 1.46.0 https://doi.org/doi:10.18129/B9.bioc.KEGGREST (2024) doi:10.18129/B9.bioc.KEGGREST.

87. Wickham, H. Ggplot2: Elegant Graphics for Data Analysis. (Springer, New York, NY, 2009). doi:10.1007/978-0-387-98141-3.

88. Kolde, R. pheatmap: Pretty Heatmaps. https://github.com/raivokolde/pheatmap (2025).

89. Perez-Riverol, Y. et al. The PRIDE database resources in 2022: a hub for mass spectrometry-based proteomics evidences. Nucleic Acids Res 50, D543–D552 (2021).

90. Deutsch, E. W. et al. The ProteomeXchange consortium at 10 years: 2023 update. Nucleic Acids Res 51, D1539–D1548 (2022).

