## Supplementary Figures for "Metagenomics-enabled proteomics reveals how AMF and PSB co-inoculation reshapes tomato rhizosphere dynamics across growth stages"

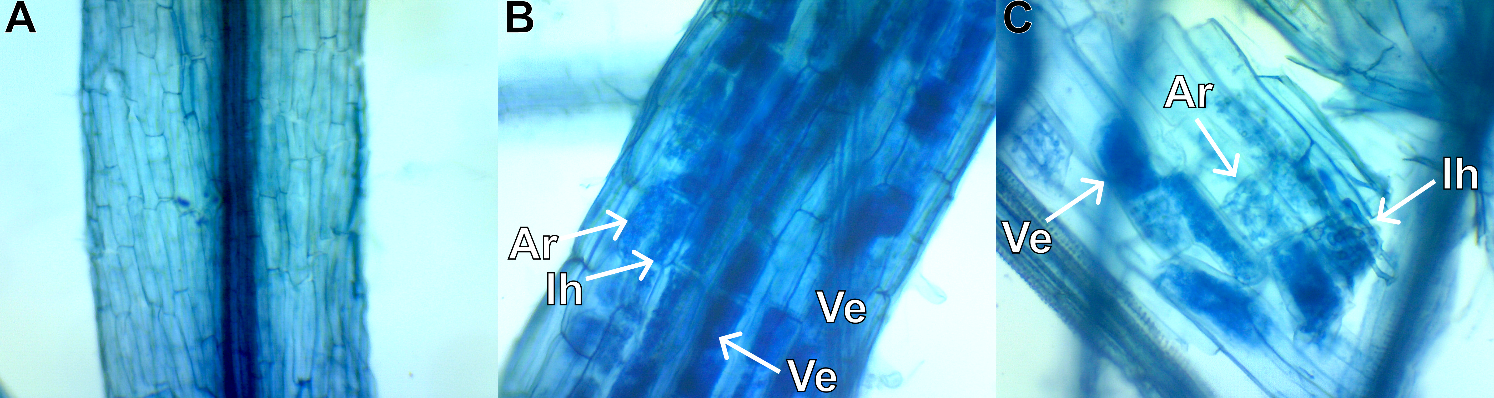

**Supplementary Figure 1.** Representative images of mycorrhizal colonization in tomato roots. (A) Uncolonized root tissue serving as a negative control. (B) High-resolution image of tomato roots heavily colonized by arbuscular mycorrhizal fungi, with clearly identifiable intraradical hyphae (Ih), arbuscules (Ar), and vesicles (Ve). (C) Magnified view of disrupted root tissue highlighting characteristic internal mycorrhizal structures.

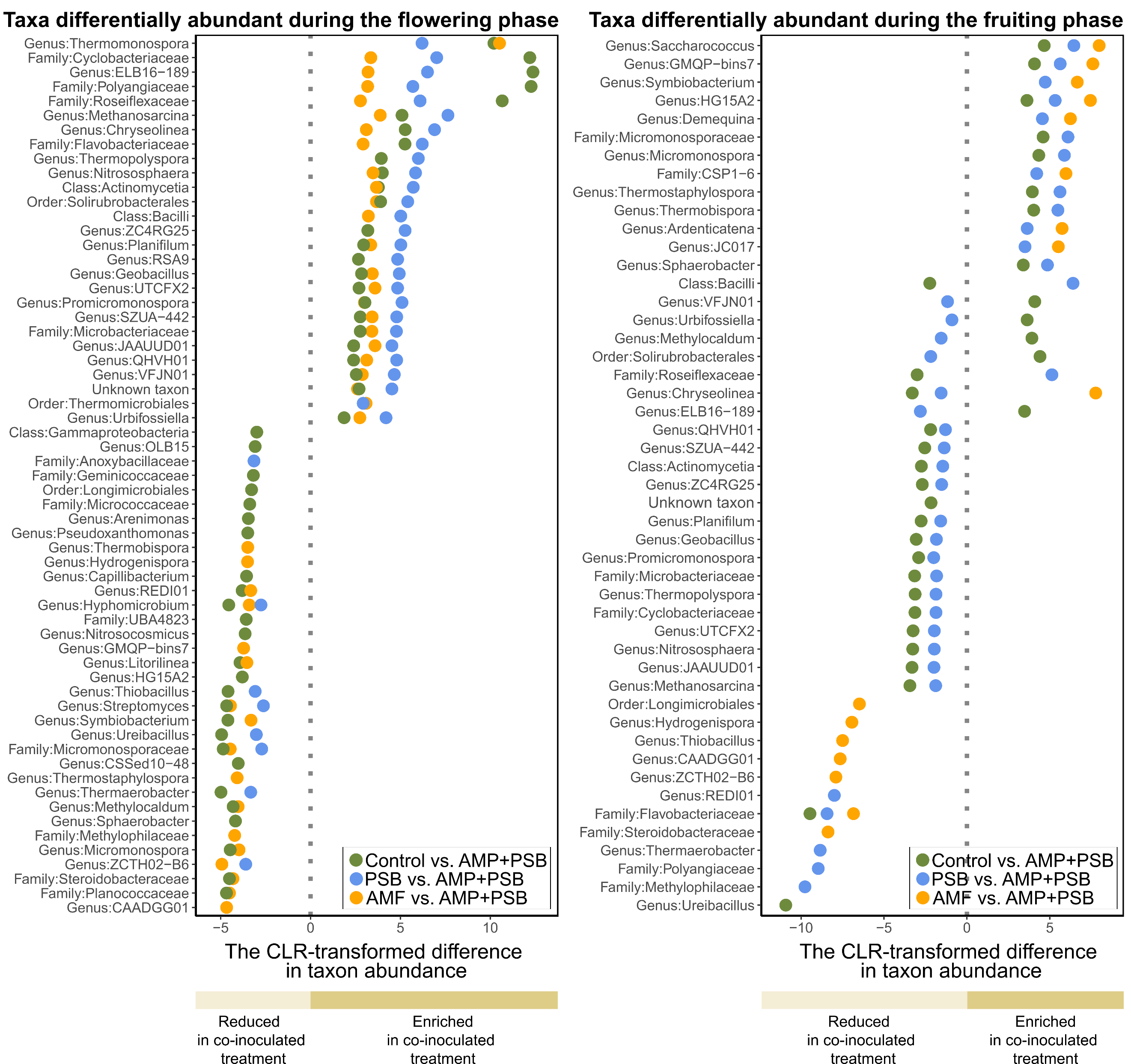

[**Supplementary Figure 2**](https://drive.google.com/drive/folders/1z3KM3uf33dSnsAI5jh6UMRptwZKp-G4Y?usp=drive_link)**.** Differential abundance analysis was performed with ALDEx2 on genus‑level 16S rRNA gene amplicon sequencing data from rhizosphere samples collected during the flowering and fruiting stages. Centered log-ratio (CLR) scores reflect shifts in taxon abundance between treatments, with positive values indicating enrichment and negative values indicating relative depletion under co-inoculation, across three pairwise comparisons. Significantly differentially abundant taxa were determined using the Wilcoxon rank-sum test with Benjamini-Hochberg correction for multiple comparisons (*p* < 0.05), and each data point is color-coded according to its respective treatment comparison. Abbreviations: Control, uninoculated tomatoes; PSB, treated with phosphate-solubilizing bacteria only; AMF, treated with arbuscular mycorrhizal fungi only; AMF+PSB, co-inoculated with both AMF and PSB.

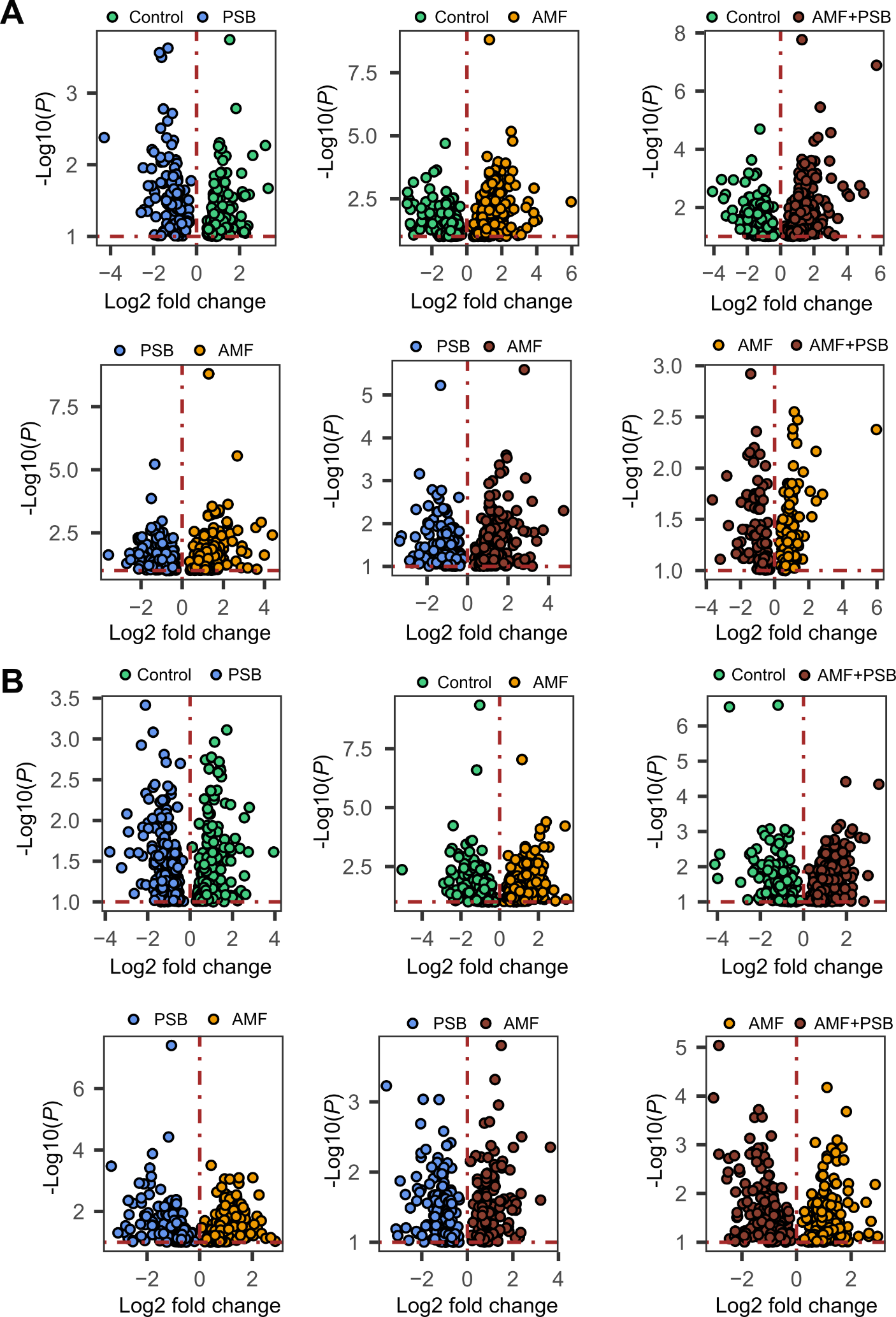

**Supplementary Figure 3.** Volcano plots illustrating differential microbial protein abundances across pairwise inoculation treatment comparisons in the rhizosphere at (A) the flowering stage and (B) the fruiting stage of tomato development. Differentially abundant proteins were identified using two-sample t-tests with Benjamini-Hochberg (BH) adjusted *p* < 0.1. The x-axis represents log2 fold change in protein abundance, with each point corresponding to an individual protein color-coded by the treatment in which it was more abundant, where the direction of fold change indicates which treatment shows greater protein enrichment in each pairwise comparison. The y-axis represents statistical significance expressed as negative log10 p-values, with higher values indicating greater confidence. A vertical dashed line at x = 0 separates proteins enriched in each compared treatment, and a horizontal dashed line marks the significance threshold (BH adjusted *p* < 0.1). Abbreviations: Control, uninoculated tomatoes; PSB, treated with phosphate-solubilizing bacteria only; AMF, treated with arbuscular mycorrhizal fungi only; AMF+PSB, co-inoculated with both AMF and PSB.

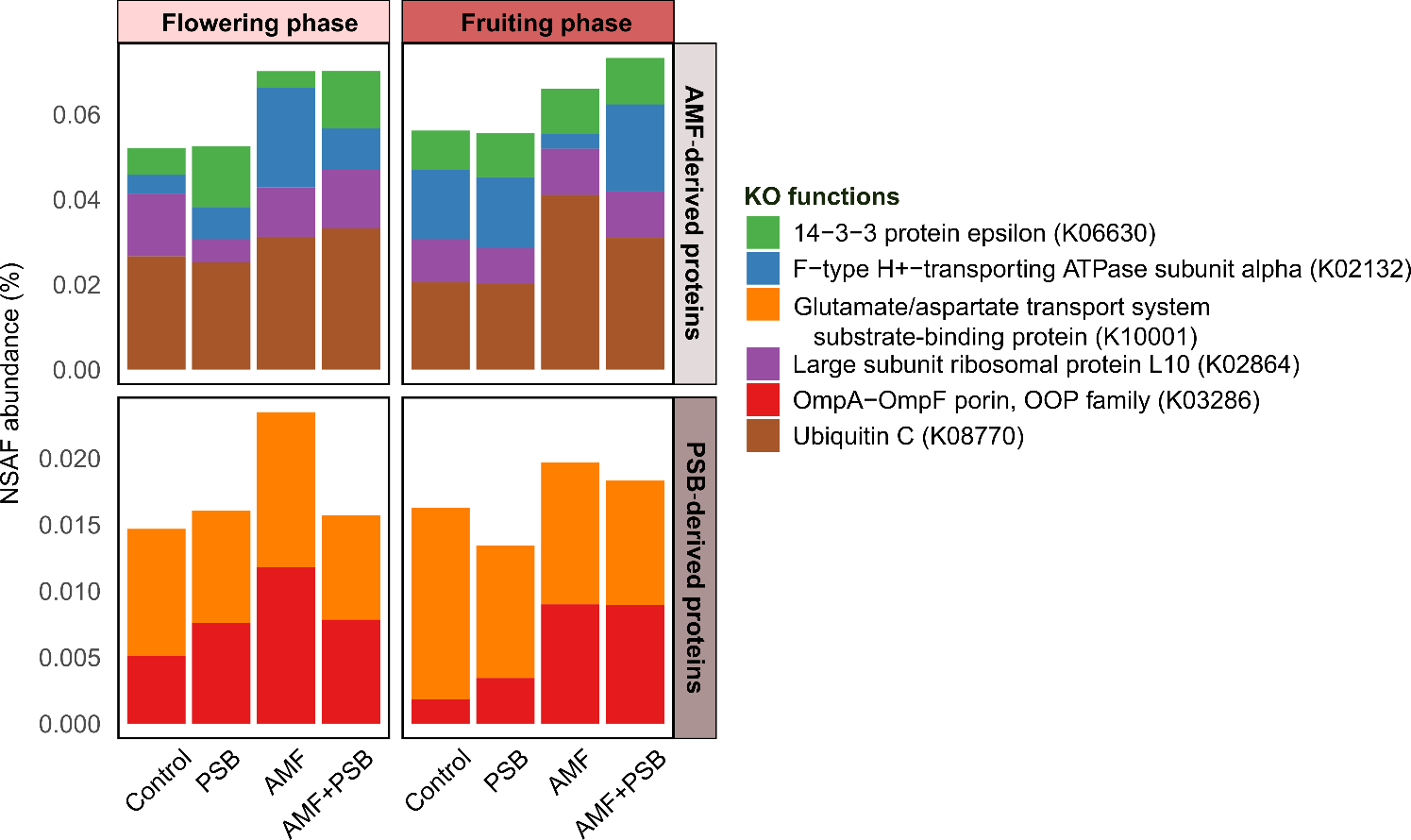

[**Supplementary Figure 4.**](https://drive.google.com/drive/folders/1z3KM3uf33dSnsAI5jh6UMRptwZKp-G4Y?usp=drive_link) Stacked bar charts depicting the mean relative abundances of proteins directly attributed to AMF inoculants (*Funneliformis mosseae* and *Rhizophagus irregularis*) and PSB inoculants (*Enterobacter cloacae* and *Pseudomonas putida*) across inoculation treatments and tomato developmental stages. Each bar segment explicitly denotes the microbial origin of detected proteins, with colors distinguishing both the taxonomic source organism and its associated functional category, enabling direct visualization of inoculant-specific proteomic contributions across treatments. Abbreviations*:* Control, uninoculated tomatoes; PSB, tomatoes treated with phosphate-solubilizing bacteria only; AMF, tomatoes treated with arbuscular mycorrhizal fungi only; AMF+PSB, tomatoes co-inoculated with both AMF and PSB.
